# A genome-resolved atlas reveals lineage-specific activity patterns of prophages in cultured human gut bacteria

**DOI:** 10.64898/2026.05.11.724261

**Authors:** Yan Huang, Ziqiang Liu, JieLin Li, Jie Hu, Xin Tan, Lei Dai, Yingfei Ma

## Abstract

Prophages dominate the gut virome, yet their basal activity landscapes remain elusive. Here, we developed ACTIVE (Activity Comparison Test for Induced Viral Elements), a robust coverage-based framework to accurately profile prophage replication activity based on sequencing data of individual bacteria. Applying ACTIVE across 31,150 gut bacterial isolates (875 species), we compiled these data into GPAC (Gut Prophage Activity Catalog) and found that 5.4% of prophages are detectably active under standard culture, with pronounced enrichment in Bacillota. Moreover, we uncovered distinct active viral lineages with broad host distributions. Comparative analyses identified 46 viral families significantly enriched for activity (SA-vFAMs). Notably, SA-vFAM7 represents a hyper-prevalent *Candidatus Heliusvirales* lineage whose host distribution expands into Bacteroidota (Tannerellaceae). Furthermore, we propose *Candidatus Mubacviridae* (SA-vFAM1), an active, hankyphage-inclusive transposable phage family associated with Bacteroidales, characterized by conserved MuA/B operons, diversity-generating retroelements, and multi-genus host distribution. Together, these findings establish GPAC as a resource linking prophage activity to host taxonomy and viral evolutionary lineages, providing a foundation for investigating prophage induction, host-phage interactions, and ecological dynamics of the human gut microbiome.

## Introduction

Prophages are pervasive and dynamic architects of the human gut microbiome^[1–3]^. By integrating into and persisting within bacterial genomes, prophages forge long-term associations that profoundly shape host fitness, interbacterial competition, and genome evolution^[4–9]^. Upon encountering specific physiological or environmental cues, these dormant elements can switch from lysogeny to active replication and lytic virion production, thereby altering microbiome structure and accelerating horizontal gene transfer^[7, 10, 11]^. Because lysogeny and prophage activity represent fundamentally different biological states, understanding the functional contribution of prophages to gut microbial ecology requires moving beyond cataloging their genomic presence to characterizing their activity^[12, 13]^. However, detectable prophage activity and its variation across human gut bacterial diversity remain insufficiently understood.

Metagenomic surveys have curated vast catalogs of gut phage sequences, exponentially expanding our view of their diversity^[14–18]^. Yet, these analyses only capture genomic insight of phages without assessing prophage activity^[14, 17, 18]^, with virus–host links further obscured by in silico predictions^[19]^. Recent efforts have been conducted to probe prophage activity. An infant cohort study observed specific activity-enriched viral clusters across populations^[20]^; an induction assay of 252 gut isolates suggested that host phylum-specific cues may govern prophage activation^[21]^. However, a systematic, host-resolved view of activity across gut prophage lineages remains lacking. The rapid expansion of large-scale cultured human gut bacterial collections offers an unprecedented opportunity to bridge this gap at single-isolate resolution^[22–30]^. De novo assemblies of tens of thousands of sequenced isolates not only anchor prophages to their definitive hosts, but their underlying raw sequencing reads also inherently capture signals of spontaneous prophage activation^[23, 26–28, 30]^. Nevertheless, unlocking this culturomics reservoir is currently hindered by the lack of a robust computational framework tailored for isolate sequencing data.

Current bioinformatic tools infer prophage activity primarily through two strategies. Alignment-based methods, such as mVIRs^[31]^ and Prophage Tracer^[32]^, detect split-reads or discordant read pairs spanning excised or circularized phage junctions. However, these approaches are sensitive to sequencing depth, confounded by abortive excision without replication^[32, 33]^, and blind to extrachromosomal prophages without classic integration^[34]^. Consequently, coverage-based approaches, exemplified by PropagAtE^[35]^, have gained traction by leveraging elevated phage-to-host read depth ratios as a direct proxy for replication-driven activity. Yet, applying PropagAtE to routine isolate sequencing data remains challenging. Its requirement for sufficient same-contig flanking host sequence to establish baseline coverage often causes it to fail in fragmented short-read isolate assemblies. In these draft genomes, prophages frequently lie near contig boundaries or span entire contigs without host context, severely truncating the repertoire of evaluable prophages and biasing cross-lineage comparisons.

Here, we developed the Activity Comparison Test for Induced Viral Elements (ACTIVE), a coverage-based, nonparametric workflow for assessing prophage activity by comparing sequencing-depth distributions between prophage regions and host genomic backgrounds. After benchmarking its performance, we applied ACTIVE to genome assemblies of cultured human gut bacteria to systematically characterize prophage activity. This analysis generated GPAC, a comprehensive, activity-annotated gut prophage atlas. By integrating activity profiles with host taxonomy and viral phylogeny, we revealed that prophage activity exhibits host- and lineage-associated patterns and identified viral lineages enriched for detectable activity. This study provides a framework and resource for investigating the distribution, evolutionary organization and ecological potential of prophage activity in the human gut microbiome.

## Results

### Development of the ACTIVE workflow for accurate prophage activity detection

To systematically profile prophage activity, we developed ACTIVE (Figure S1a). This approach rests on the premise that active prophage replication increases viral DNA copy number, thereby elevating sequencing depth across the prophage region relative to the host chromosome^[35, 36]^. ACTIVE quantifies this relative coverage enrichment using a statistical comparison and Cliff’s Delta, a robust effect size quantifying the separation between prophage and host-background depth distributions, to operationally classify prophages as active^[37]^ (Figure 1a). To accommodate differences in genome completeness and assembly quality, ACTIVE implements two modes for estimating host background coverage: a *’Fast*’ mode that uses host-conserved single-copy marker genes (MGs)^[38]^; and a ’*Precise*’ mode that uses the entire non-phage portion of the genome (Figure 1a).

**Figure 1.**
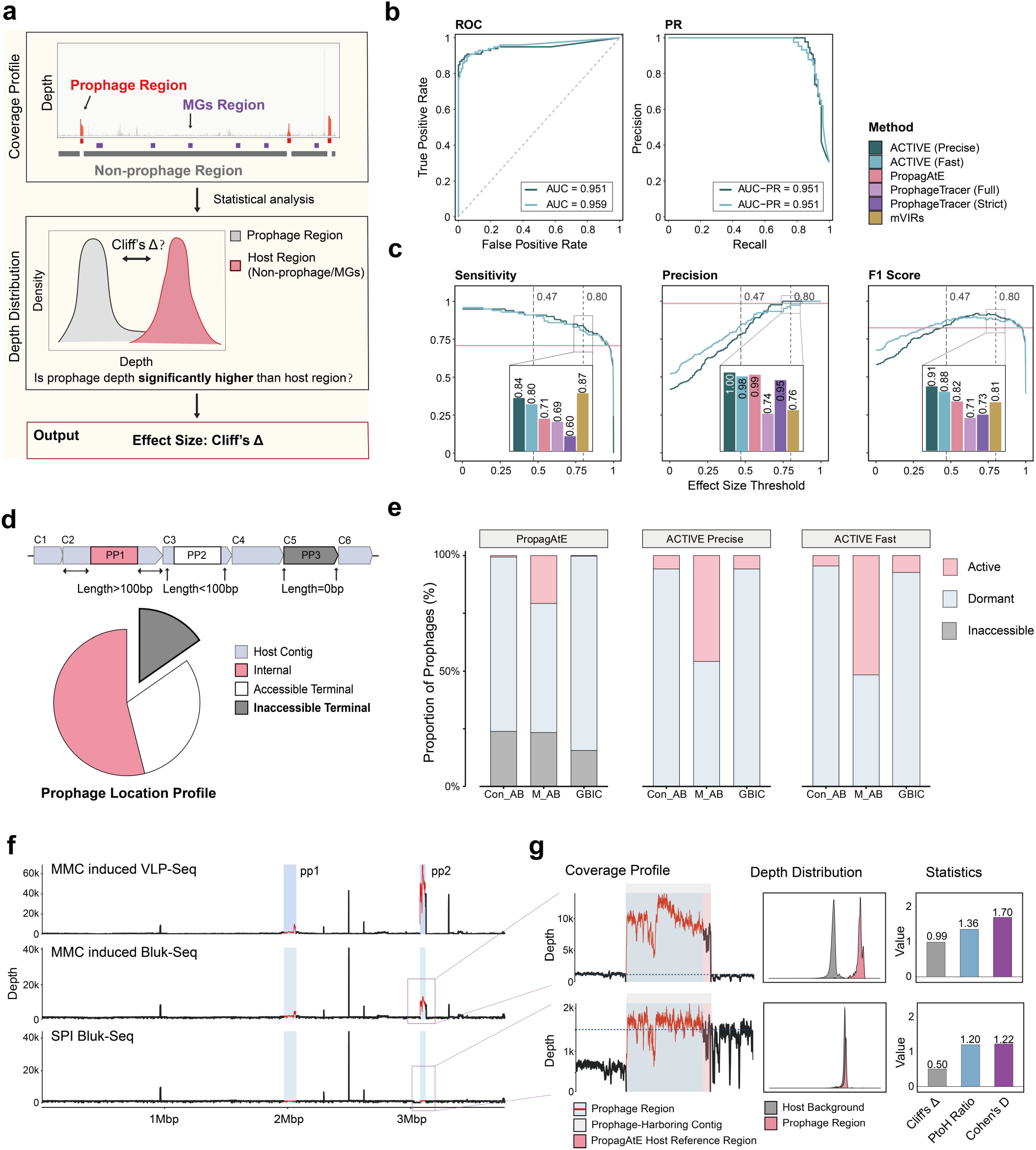
Evaluation of Prophage Activity via ACTIVE. (a) Schematic of the ACTIVE framework. ACTIVE evaluates prophage activity by comparing prophage and host depth distributions using Cliff’s Delta (active: ≥ 0.8). Host regions are defined as flanking MG regions (Fast mode) or non-prophage genomic regions (Precise mode). (b) ROC and PR curves obtained by varying the Cliff’s Delta threshold (Benchmark 1). Colors denote the ACTIVE mode. (c) Benchmarking across Cliff’s Delta thresholds (Benchmark 1). Curves show sensitivity, precision, and F1 score. Dashed lines mark the large-effect (0.47) and empirical (0.8) thresholds; pink lines indicate PropagAtE performance. Boxed bars compare tools at Cliff’s Delta = 0.8 (Prophage Tracer shown under Full and Strict criteria, Methods). (d) Prophage configurations in fragmented genomes. Left: Schematic of internal (PP1), accessible-terminal (PP2), and inaccessible-terminal (PP3) prophages based on distance to contig ends. The chart shows the prophage location profile across the A. baumannii and GBIC datasets. (e) Performance comparison on the *A. baumannii* (AB) and GBIC datasets. ACTIVE classifies prophages as active (Cliff’s Delta ≥ 0.8) and dormant (Cliff’s Delta < 0.8). Gray denotes terminal prophages inaccessible to PropagAtE. Con_AB means *A. baumannii* under SPI, M_AB means *A. baumannii* under MMC induction. (f) Whole-genome depth profiles of strain RAB20B under spontaneous induction (SPI), mitomycin C (MMC)-induced bulk sequencing, and MMC-induced VLP sequencing. Predicted prophage regions are indicated by red lines and blue shading. (g) Local coverage profiles of prophage pp2 in strain RAB20B (±30 kb) under SPI and MMC induction. Blue/red: prophage; gray: prophage-containing contig; pink: PropagAtE host reference. Right panels display ACTIVE depth distributions (Precise mode) and activity metrics (PtoH (Prophage to Host ratio), Cliff’s Delta, and Cohen’s d). PropagAtE classifies a prophage as active when Cohen’s d ≥ 0.7 and PtoH ≥ 2, whereas ACTIVE uses Cliff’s Delta ≥ 0.8.

We first benchmarked ACTIVE using a reference dataset comprising 348 experimentally characterized prophage induction events across multiple bacterial species. The dataset included complete host genomes, curated prophage coordinates, and matched bulk sequencing datasets generated under various induction conditions^[31, 32, 35, 39–43]^ (Figure S1c, Benchmark 1; Table S1). We compared ACTIVE with three existing approaches: the coverage-based tool PropagAtE, the split-read/clipped-alignment-based methods Prophage Tracer and mVIRs. Receiver-operating-characteristic (ROC) and precision–recall (PR) analyses showed that Cliff’s Delta effectively discriminated experimentally characterized activity states (Figure 1b). Beyond the conventional large-effect threshold (≥ 0.47)^[44]^, ACTIVE outperformed PropagAtE in both sensitivity and F1-score (Figure 1c). At Cliff’s Delta ≥ 0.8, ACTIVE reached peak precision while maintaining high sensitivity, achieving the highest overall F1-score among all evaluated methods (Figure 1c, S1d). Based on these results, we operationally classified prophages as active when they exhibited statistically significant coverage enrichment and a Cliff’s Delta of ≥ 0.8, as it falls within the F1-score plateau in the benchmark and provides stringent discrimination between prophage and host-background coverage distributions (Figure S1e, S2b). We further evaluated the robustness of ACTIVE using short-read assemblies reconstructed from the Benchmark 1 datasets (Figure S1c, Benchmark 2). Despite assembly fragmentation, with 48.2% of prophages spanning multiple contigs, the *Precise* mode retained strong performance on assembled genomes (F1-score = 0.88; Figure S1f-h; Table S2).

We next evaluated ACTIVE on two independent isolate collections comprising 63 *Acinetobacter baumannii* isolates and 442 gut bacterial isolates representing diverse species (Table S3-S5). Across these datasets, the *Fast* and *Precise* modes yielded highly concordant Cliff’s Delta estimates (Spearman’s r = 0.927, P < 0.001; Figure S2c), indicating that the two modes provide consistent activity assessments despite differences in how host-background coverage is defined. We then examined the extent to which genome fragmentation affects prophage activity assessment. Among 962 predicted prophages in these isolate datasets, 443 (46.0%) were located within 100 bp of a contig boundary and were therefore classified as terminal prophages (Figure 1d). Of these, 35.0% (155/443) lacked host-flanking sequence and could not be evaluated by PropagAtE (Figure 1d, inaccessible terminal; Table S5, host_len = 0). In contrast, ACTIVE could assess such prophages by using non-prophage genomic regions or host-conserved marker genes as the host background, without requiring host-flanking sequence on the same contig (Figure 1e). A similar limitation of PropagAtE was observed in Benchmark 2, where approximately 53% of prophages could not be evaluated (Figure S1g). Thus, in addition to its benchmarked classification performance, ACTIVE expands the computable range of prophage activity inference in fragmented isolate assemblies.

To experimentally validate ACTIVE predictions, we performed paired bulk and virus-like particle (VLP) sequencing, together with transmission electron microscopy (TEM) on samples from selected *A. baumannii* isolates under spontaneous induction (SPI) and mitomycin C (MMC)-induction conditions (Figure 1f-g, S2d-e, Table S6). In strains RAB20B and SAB43B, VLP-associated DNA yields were insufficient for library construction under spontaneous induction with no active prophage predicted, whereas MMC induction generated sufficient VLP-associated DNA. In the MMC-induced sample, prophage regions identified as active by ACTIVE displayed pronounced, prophage-specific coverage enrichment in VLP-seq data, and TEM confirmed the presence of tailed phage particles in supernatants (Figure 1f-g, S2b). These experimental results resolved discrepancies between PropagAtE and ACTIVE. In strain RAB20B, ACTIVE identified prophage pp2 as active, whereas PropagAtE classified it as dormant because short host-flanking regions provided an unrepresentative local coverage baseline (Figure 1g). Similarly, in strain SAB43B, ACTIVE detected weak but significant coverage enrichment in prophage regions, whereas PropagAtE classified pp1 as dormant (Figure S2d). Notably, although split-read analysis failed to detect the activity of prophages pp2 and pp3, it partitioned pp1 into a PICI-like element^[45]^ and an adjacent prophage (Figure S2g, Table S4). In line with this, coverage profiling across pp1 revealed two distinct peaks that were clearly separated from the host background (Figure S2e).

Together, these results demonstrate that ACTIVE provides a robust framework for assessing prophage activity across fragmented bacterial genome assemblies and can identify prophage regions exhibiting replication-associated coverage enrichment that may be missed by existing approaches. Experimental validation further supports the activity predictions for selected *A. baumannii* isolates. These findings provide a basis for applying ACTIVE to large-scale, genome-resolved analyses of cultured bacterial isolates.

### GPAC establishes a culturome-derived, activity-annotated atlas of human gut prophages

To comprehensively characterize prophage activity across human gut bacteria, we analyzed 31,150 quality-controlled genome assemblies from five publicly available cultured gut bacterial isolate collections^[26–28, 30]^ during standard *in vitro* growth conditions, spanning 875 species across 12 phyla (Figure 2a, Table S7). By integrating a multi-tool prediction and quality-curation pipeline (Figure S2a), we compiled 54,160 high-confidence prophages into the Gut Prophage Activity Catalog (GPAC), establishing a culturome-derived, activity-annotated atlas (Figure S3a). The collection included terminal prophages that are inaccessible to PropagAtE, thereby extending activity profiling to prophages in fragmented isolate assemblies (Figure S3b).

**Figure 2.**
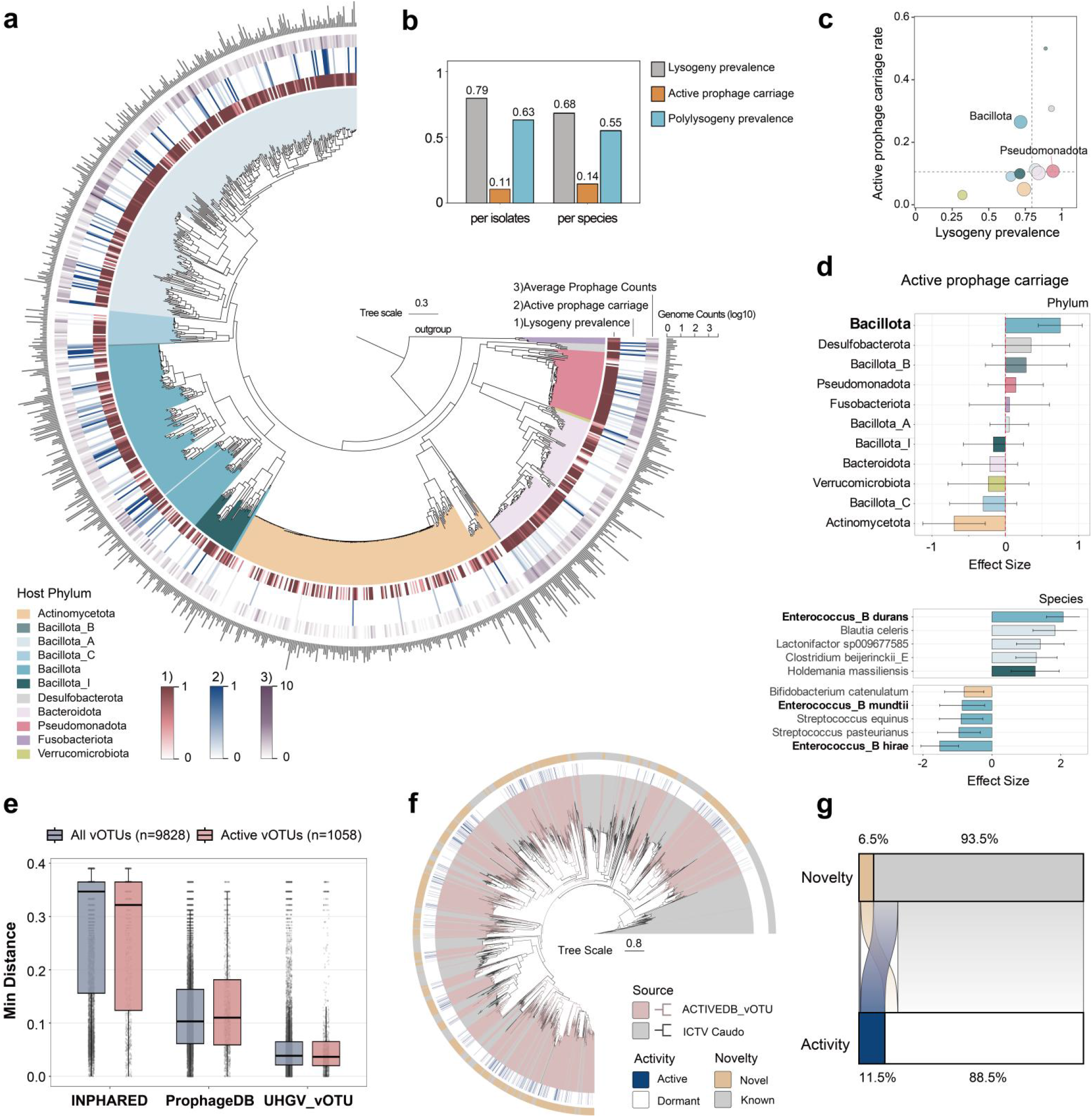
Construction and characterization of GPAC. (a) Phylogenetic tree constructed from GTDB-Tk-derived conserved marker genes. Rings from inside to outside: host phylum; lysogeny prevalence (lysogens/total strains); active prophage carriage (lysogens carried active prophage / total lysogens); average prophage count per lysogen and number of genomes per species (bar plot, log_10_). (b) Descriptive statistics of host prophage-associated traits. Lysogeny prevalence and active prophage carriage are mentioned as above. Polylysogeny prevalence means the fraction of lysogens with >1 prophage. "Per species" denotes weighted averages of species-level rates that account for uneven genomic sampling across different species (Methods). (c) Phylum-level summary of lysogeny prevalence and active prophage carriage. Dots are sized by phylum membership and colored by phylum. (d) Taxonomic rankings of active prophage carriage as predicted by GLMM. Effect sizes on the X-axis represent condition-adjusted marginal deviations from the baseline (vertical red dashed line = 0). Error bars define the conditional standard errors of the model estimates. (e) Minimum genomic distances between GPAC prophages and reference databases (INPHARED, ProphageDB, and UHGV), stratified by all vOTU and active vOTU (AvOTUs). (f) TerL-based phylogenetic tree of *Caudoviricetes* vOTU in GPAC. Branch colors distinguish ICTV genomes from GPAC vOTUs. The inner ring marks vOTU activity (Active: containing ≥1 active genome; Dormant: all genomes inactive). The outer ring indicates novelty (Known: assigned to ICTV ranks from species to order; Novel: unassigned). (g) Sankey diagram connects the activity state with the taxonomic novelty of each *Caudoviricetes* vOTU.

GPAC revealed that lysogeny was pervasive across the human gut microbiota, occurring in 79.4% of isolates, with 63.0% of lysogens (15,593/24,746) harboring multiple prophages (Figure 2b). Within this lysogenic reservoir, ACTIVE identified 2,920 active prophages spanning 2,610 lysogenic strains across 242 species and 11 distinct phyla. These findings establish an *in vitro* activity baseline consistent with spontaneous induction (5.4% active; median mPtoH = 1.98 (IQR: 1.69–2.80); Figure S3d). Active prophage carriage was distinctly structured by host taxonomy. Whereas Pseudomonadota and Bacteroidota exhibited high lysogeny prevalence but low frequencies of active prophage carriage, Bacillota was enriched in both traits (Figure 2c). Hierarchical generalized linear mixed models (GLMMs) controlling for genome assembly quality and isolation source confirmed this pattern, identifying Bacillota as the phylum most strongly associated with active prophage carriage (Figure 2d, phylum panel; Table S8; Methods). This positions Bacillota as the primary phylum-level reservoir of active prophages activity. Variance partitioning further indicated substantial contributions of genus- and species-level effects to prophage-associated traits, even within Bacillota (Figure S3f). For example, *Enterococcus_B durans* exhibited the strongest positive association with active prophage carriage, whereas its relative *Enterococcus_B hirae* displayed the strongest negative association (Figure 2d). More broadly, closely related species frequently displayed divergent lysogeny–activity traits across bacterial phyla (Figure S3g). These findings demonstrate that while active prophages are widely distributed across the host phylogeny, their distribution exhibits pronounced host-lineage-specific variation down to the species level, with Bacillota serving as the predominant reservoir.

Beyond host-level patterns, GPAC uncovers extensive viral diversity and resolves previously uncharted active lineages. Genomic and proteomic clustering resolved the collection into 9,828 viral operational taxonomic units (vOTUs), 3,976 viral genera (vGENUS) and 305 viral families (vFAMs) (Table S9), highlighting the extensive diversity of the collection. Focusing on this active fraction, we applied previously established criteria^[21]^ and identified 1,058 vOTUs (10.8%) as active vOTUs (AvOTUs), each containing at least one active prophage. Genomic comparisons revealed that nearly half of these active prophages were absent from ICTV or UHGV references^[16]^ (Figures S3h–i). Importantly, GPAC bridges the gap between isolated prophages and uncultivated metagenomic dark matter, displaying genomic distances closer to the uncultivated gut virome (UHGV) than to classical cultured databases (INPHARED^[46]^) (Figure 2e). Since Pharokka^[47]^ annotated terminase large subunit (TerL) in 77% of vOTUs (Figure S3j), we performed phylogenetic profiling on the 6,711 Caudoviricetes vOTUs encoding a single TerL. This analysis revealed deep taxonomic novelty, with 93.5% unclassified below the class level, coalescing into numerous expansive, ICTV-underrepresented subclades (Figure 2f-g). Remarkably, 11.5% of these *Caudoviricetes* lineages represented AvOTUs, the vast majority of which define previously uncharacterized clades (Figure 2g). Together, these findings establish GPAC as a resource that annotates prophage activity across cultured gut bacteria, reveals host-lineage-specific activity patterns and substantial novel viral diversity.

### Active prophages exhibit diverse lineage origins and broad cross-host distribution

To resolve prophage activity at the viral species level, we mapped the activation profiles across all 1,058 AvOTUs in GPAC. The proportion of active members varied considerably among AvOTUs, suggesting distinct, viral lineage-specific activity propensities (Figure 3a). Capitalizing on this heterogeneity to pinpoint significantly activity-enriched gut viral lineages while controlling for culture-source variations, we applied a source-stratified exact test framework to AvOTUs containing ≥ 5 member genomes (Methods). Taxa with significantly higher active proportion compared to background levels (FDR < 0.05, common OR > 1) were defined as significantly activity-enriched vOTUs (SA-vOTUs) (Table S10). In total, we identified 94 SA-vOTUs, the largest fraction of which was associated with Bacillota (n=43) (Figure 3a and S4a). Notably, SA-vOTU4777 was the most statistically prominent taxon (110/192, Padj = 1.36×10^-76^, OR = 20.91) (Figure 3a). This SA-vOTU was identified as a P4-like phage satellite carried by *Escherichia coli*^[45]^ (Figure S4c). Accumulation curve analysis showed that the discovery of both vOTUs and AvOTUs remains far from saturated, highlighting extensive unexplored diversity, whereas SA-vOTUs discovery reached a plateau, suggesting that the core active viral lineages are largely captured (Figure 3b).

**Figure 3.**
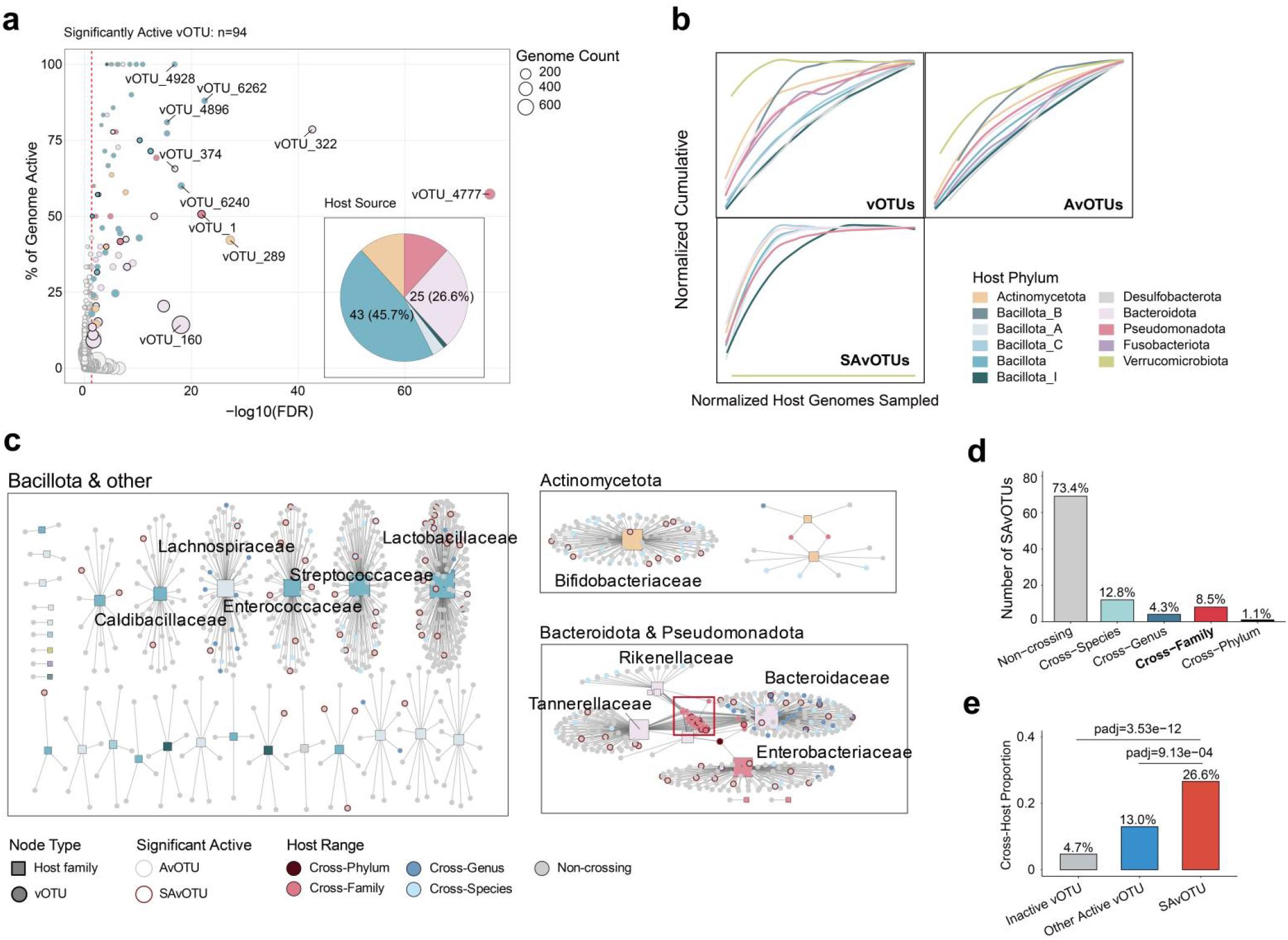
Active prophages with broad host distribution. (a) Activity landscape of AvOTUs. The x-axis represents statistical significance (−log₁₀ (FDR), Benjamini-Hochberg corrected), and the y-axis indicates the prevalence of activity (% genome active) for each vOTU. Non-significantly active vOTUs: grey. Colored: SA-vOTUs, colored by host phylum. Bold: active vOTUs distributing different species (cross-host). Node size: number of constituent genomes. Red dashed line: significance threshold (FDR = 0.05). (b) Accumulation curves of vOTUs, AvOTUs, SA-vOTUs across host phyla. (c) Distribution network between active vOTUs and host families. The top 10 highly connected host families and top 10 most significant SAvOTUs are labeled. Colored: Cross-host active vOTUs. Bold: SA-vOTUs. (d) Composition of SA-vOTUs by cross-host distribution degree. Dormant vOTU: vOTU with no active member. Other active vOTU: AvOTU not reaching Fisher’s exact test significance. (e) Comparison of the proportion of cross-host vOTUs (defined as cross-species or higher) among different activity categories. FDR derived from Fisher’s exact test.

Given that AvOTUs were distributed across diverse host phyla (Figure 3b), we constructed a host-virus interaction network to characterize the host distribution of AvOTUs. AvOTUs were enriched in Bacteroidota, Bacillota (specifically Enterococcaceae, Streptococcaceae, and Lactobacillaceae) and Enterobacteriaceae (Figure 3c). Analysis of an integrated host-association map across all vOTUs further revealed major clusters enriched for cross-host associations, consistent with metagenomic predictions for Lachnospiraceae and Bacteroidales^[1, 14]^ (Figure 3c and S3k). These results delineate the primary host reservoirs of AvOTUs and reveal their broad host distribution. Overall, nine SA-vOTUs (9.6% of all SA-vOTUs) had exceptionally broad host distributions, spanning host families or higher taxonomic ranks (Figure 3d). Importantly, cross-host distributions were significantly more frequent among SA-vOTUs (Figure 3e). These SA-vOTUs were primarily distributed across Bacteroidota and served as key nodes for cross-family interaction (Figure 3c). Among them, SA-vOTU4771 (57/279, Padj = 1.11×10^-17^, OR=4.60) had the broadest cross-host distributions, bridging Bacteroidota and Pseudomonadota (Figures 3c, S4d-e). Together, these findings identify a core assemblage of active gut prophage species. Their significant active proportion and putative broad host distribution mark them as likely major contributors to virus-host dynamics across the human gut microbiome.

### Family-level analysis defines active and exceptionally prevalent viral lineage

While individual vOTUs capture extensive diversity, vFAMs provide a higher-level framework for identifying dominant gut viral lineages^[48, 49]^, for which discovery approached saturation in our dataset (Figure S4f). To address whether activity is enriched in specific viral lineages, we defined active vFAMs (AvFAMs) as vFAMs containing at least one active member. By this criterion, ∼54% of vFAMs (166/305) were classified as AvFAMs. Applying the same source-stratified statistical framework to the viral family level (≥ 5 members per AvFAM), we identified 46 significantly activity-enriched vFAMs (SA-vFAMs; FDR < 0.05, common OR > 1) (Figure 4a and S4d, Table S10), indicating lineage-specific activity. Most SA-vFAMs were associated with Bacillota, with a mean proportion of active members exceeding 30% (Figure 4b). Ten SA-vFAMs were hyper-active (with more than 50% of their members active), and eight of these (80%) were associated with Bacillota (Table S10). This phylum-level enrichment highlights Bacillota as a primary reservoir of active gut prophages. Several SA-vFAMs contained exceptionally large numbers of members, suggesting extensive diversification in the gut. Notably, four of the ten largest SA-vFAMs lacked ICTV classification below the class *Caudoviricetes* level (Figure S5a), highlighting the extent of unclassified diversity within active gut lineages. These included SA-vFAM9 (149/1,524, Padj = 4.42 × 10⁻ ¹², OR = 2.03), a Bacteroidales-associated family characterized by significantly shorter genome lengths in active members compared to dormant counterparts (Figure S5a-d).

**Figure 4.**
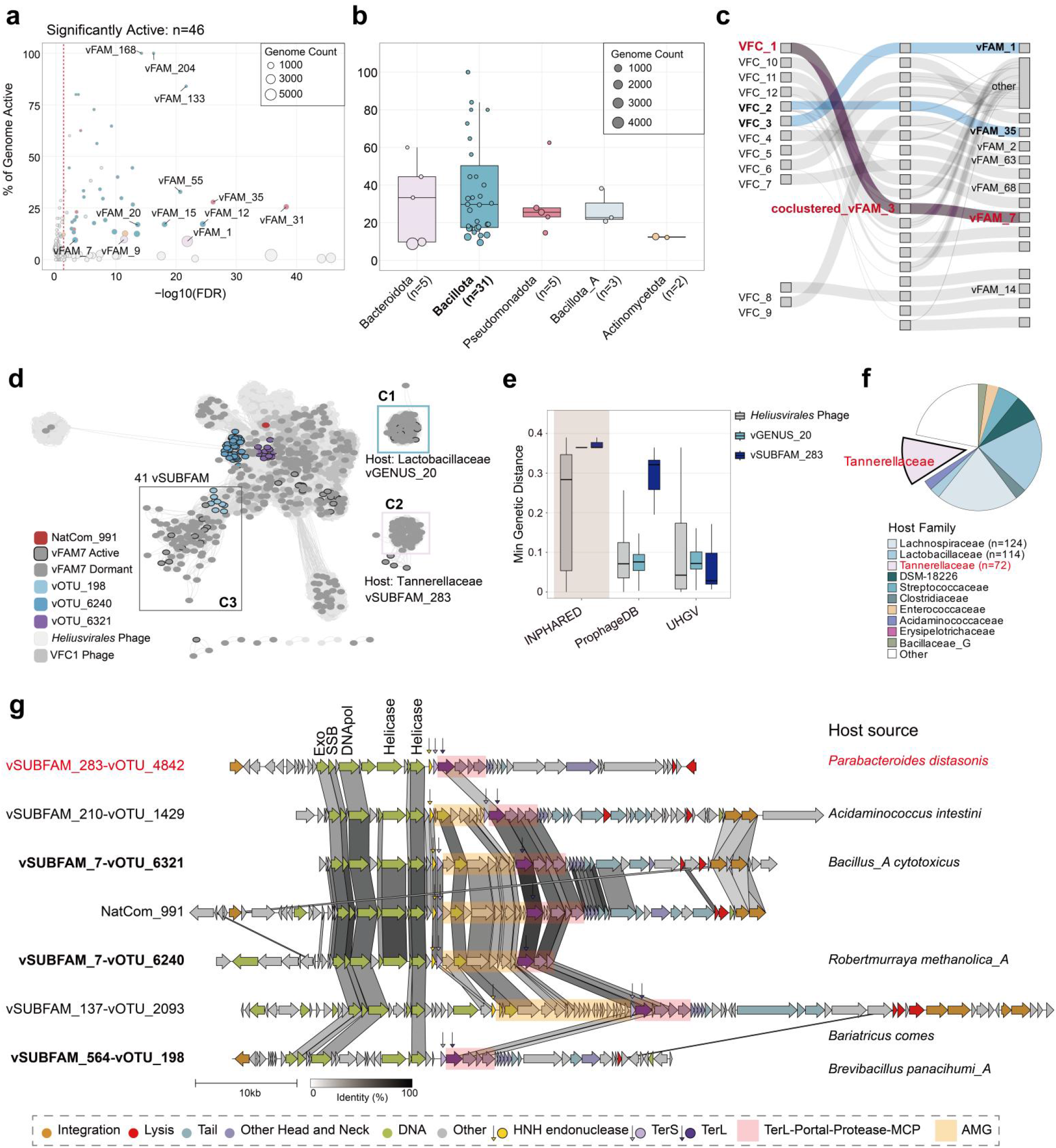
A SA-vFAM with High Prevalence. (a) Activity landscape of all AvFAMs illustrates the percentage of active genomes (y-axis) and statistical significance (x-axis, -log_10_(FDR), FDR derived from Fisher’s exact test (Benjamini-Hochberg)). Non-significant AvFAMs are shown in grey. SA-vFAMs are colored according to their respective host phyla. Red dashed line: significance threshold (FDR = 0.05). (b) Proportion of active members of SA-vFAMs across various host phyla. (c) Co-clustering of SA-vFAMs with high-prevalence VFCs identifies both highly prevalent and active viral families. Sankey diagram illustrating the co-clustering patterns between VFCs and vFAMs. VFCs co-clustering within SA-vOTUs are colored, and those forming complete co-clusters with SA-vFAMs are highlighted in dark blue. Similarly, SA-vFAMs co-clustering with VFCs are colored and annotated, with complete co-clusters also marked in dark blue. The specific co-clustering path comprising VFC_1, NvFAM3, and vFAM7 is highlighted in purple. (d) Proteomic similarity network between VFC_1 and 1,032 *Ca. Heliusvirales* reference genomes, generated using vConTACT2. The three newly identified viral lineages are outlined in boxes. (e) Minimum genomic distances between SA-vFAM7 and reference databases (INPHARED, ProphageDB, and UHGV), stratified by *Ca. Heliusvirales,* vGENUS20 and vSUBFAM283. (f) Family-level host distribution of vFAM7. (g) Comparative genomics and consensus host taxonomy for the six largest vFAM7 subfamilies. AMGs and the conserved head–neck modules are highlighted.

We next asked whether these active viral families correspond to globally prevalent lineages by co-clustering our vFAMs with 12 reference high-prevalence viral family clusters (VFCs; detected in >50% of over 3,000 global human gut virome samples)^[50]^ (Methods). Strikingly, several SA-vFAMs co-segregated almost exclusively with specific VFCs, notably SA-vFAM7 with VFC_1 (Figure 4c, Table S11). Because VFC_1 represents the most prevalent lineages across human populations (> 90%)^[50]^, this co-clustering suggests vFAM7 as both active and globally prevalent gut phage lineage (Figure 4c, Table S11). SA-vFAM7 was a member-rich, diverse and significantly activity-enriched family (9.48% of its 591 prophage members active; Padj = 6 × 10^-4^, OR = 1.82), with 3 SA-vOTUs among its 279 vOTUs, indicating within-family lineage specificity (Figure S5e). We therefore characterized SA-vFAM7 in depth.

SA-vFAM7 was placed within both *Ca. Heliusvirales*^[51]^ and VFC_1 by TerL phylogeny and proteomic-sharing network (Figures 4d, S5f), and resolved into three sublineages: the Lachnospiraceae-associated vGENUS20 cluster (C1), the Tannerellaceae-associated vSUBFAM283 cluster (C2), and a highly diverse, loosely clustered group comprising 41 vSUBFAMs (C3). This substructure alone indicates unexplored diversity within *Ca. Heliusvirales*, at least at the subfamily level. Two of these sublineages are notable for their novelty and host breadth: C1 and C2 showed no sequence homology to INPHARED yet were closely related to uncultivated UHGV genomes (Figure 4e), and C2 extended the known host distribution of *Ca. Heliusvirales* from Bacillota to Bacteroidota^[51]^, indicating unexplored ecological versatility (Figure 4f). Despite this diversification, SA-vFAM7 retained the core genomic architecture of *Ca. Heliusvirales*, including five early replication proteins and a TerL–portal–proteinase–major capsid protein (MCP) structural module^[51]^ (Figure 4g). Specific clades (vSUBFAM7) harbored AMGs within the packaging module (defined here as HNH endonuclease – TerS – TerL), with variable insertion sites and content (Figure S5g-h). In summary, family-level profiling characterizes SA-vFAM7 as an active, diverse, and globally prevalent *Ca. Heliusvirales* lineage with an expanded host distribution.

### A novel Mu-like transposable phage family, *Candidatus Mubacviridae*, shows high activity and broad host distribution

To investigate the genetic features associated with prophage activity, we analyzed high-quality Caudoviricetes genomes (completeness ≥ 90%) in GPAC. As expected, active prophages preferentially encoded structural components, whereas dormant prophages were enriched for AMGs and DNA metabolism genes (Figure S6a)^[21]^. Unexpectedly, however, transposases and their associated domains were ranked among the most enriched features in vOTUs with high proportions of active genomes (Figure 5a, Table S12). Targeted synteny analysis revealed that transposases frequently formed an adjacent, co-directional operon with the DNA transposition protein MuB (Figure 5b–c, Table S13), a hallmark of transposable Mu-like phages. MuA/B-encoding prophages indeed showed a higher proportion of active genomes than other prophages, indicating that Mu-like prophages, defined by MuA transposase and MuB AAA ATPase^[21, 52]^, are both widespread and preferentially active in GPAC (Figure S6b). Screening for vFAMs dominated by MuA/B phages identified six vFAMs, four of which were active and primarily associated with Enterobacteriaceae and Bacteroidales hosts (Figure S6c-d). These vFAMs possessed complete structural modules aligned with ICTV-defined Mu-like prophages^[53, 54]^. Three of them represented taxonomically novel families associated with hosts in the order Enterobacterales (vFAM257, vFAM152 and vFAM89; Figure S6e). Comparative genomic placement anchored these newly identified Mu-like vFAMs within the global diversity of transposable phages, revealing that they represent previously uncultured, gut-associated lineages, even in well-studied hosts such as Escherichia coli. Accordingly, these vFAMs showed close genomic similarity to metagenomic genomes in the Transposable Bacteriophage Genome Database (TBPGD)^[55]^ and UHGV, yet marked divergence from isolated reference phages in INPHARED (Figure 5d).

**Figure 5.**
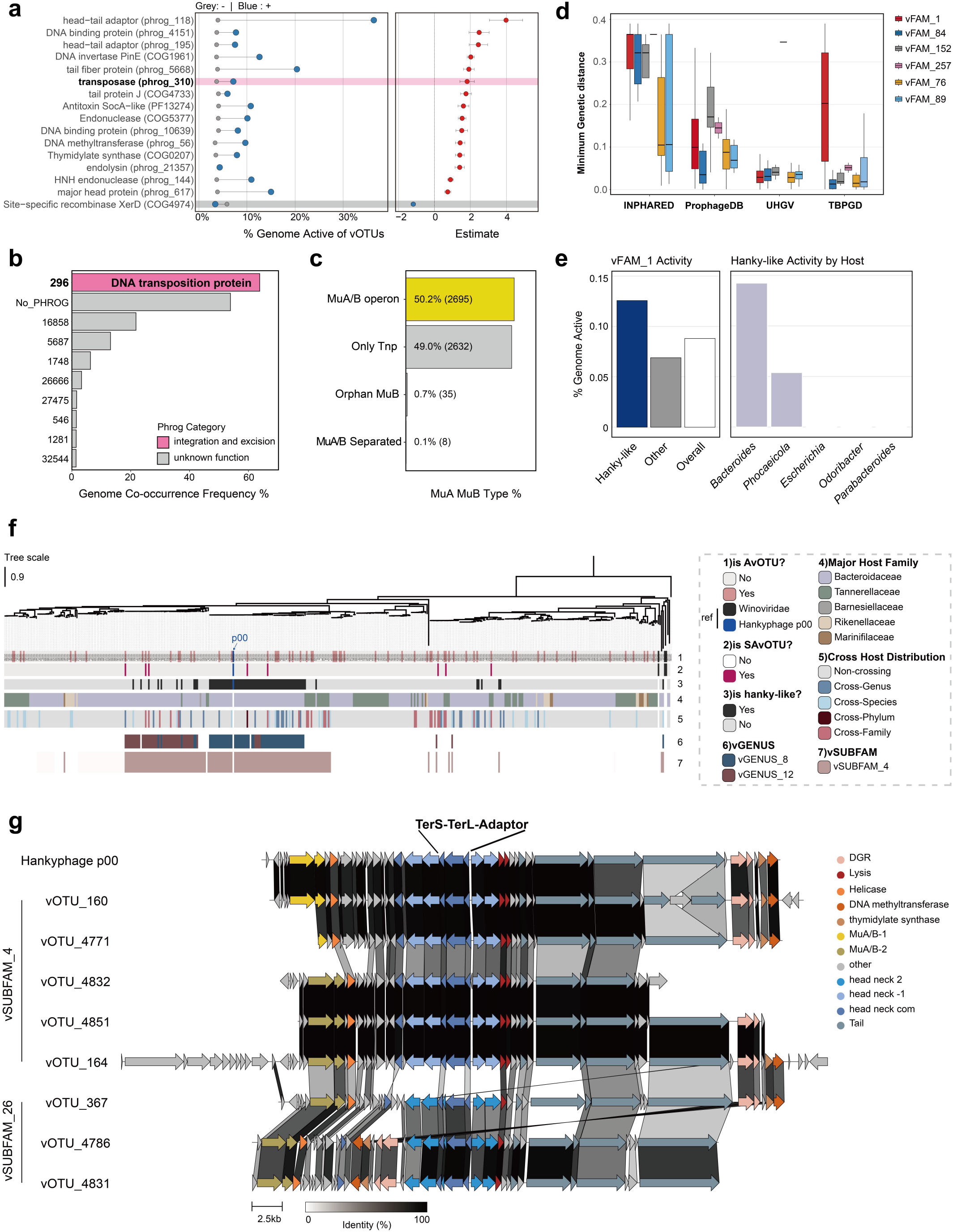
Genomic Feature Enrichment Identifies a Highly Active Family *Candidatus Mubacviridae* (SA-vFAM1). (a) Top 10 core protein domains positively correlating with prophage induction, detailing the differences in proportion of active prophages and their corresponding GLMM-derived statistical significance. (b) Top 10 transposase-adjacent genes (±1 CDS, same transcriptional orientation) identified across transposase-encoding *Caudoviricetes*, categorized by PHROG and functional category. Frequencies indicate the percentage of genomes harboring each specific gene. (c) Co-occurrence patterns of transposases and MuB (COG2842) across *Caudoviricetes*. MuA/B operon: adjacent and co-directional transposase and MuB. MuA/B separated: co-occurring but non-adjacent. Only Tnp: transposase present without MuB. Orphan MuB: MuB present without transposase. (d) Minimum genomic distances between the six Mu-like phage vFAMs and the reference databases INPHARED, ProphageDB, UHGV, and TBPGD. Comparisons with TBPGD were based on all genomes within each vFAM, whereas the other comparisons were based on the vOTU representative genomes within each vFAM. (e) Quantitative assessment of proportion of active prophages. Bar plots detail (i) the global activity comparison among hankyphage members, non-hankyphage members, and SA-vFAM1 overall (ii) the host genus-specific activity exclusively for identified hankyphages. (f) A TerL-based vOTU tree of SA-vFAM1. Rings from inside to outside represent: (1) AvOTU (colored); (2) SA-vOTU (colored); (3) Detected hanky-like prophages in vOTUs (in black); (4) Major host family; (5) Cross-host distribution; (6) vGENUS of vOTUs; (7) vSUBFAM of vOTUs. The arrow marks the hankyphage reference strain p00. (g) Comparative genomic architecture of representative SA-vOTUs within SA-vFAM1. Emphasizing the robust conservation of the TerS-TerL-Adaptor packaging module.

Among these six MuA/B-dominated vFAMs, SA-vFAM1 (8.81% active genomes; 404/4,584; Figure 5e) stood out prominently. It represents the largest family among all SA-vFAMs in GPAC (Figure 4b), encompassing the vast majority (82.3%, 2,132/2,589) of all detected MuA/B prophages (Figure S6b). Ecologically, SA-vFAM1 operated as a multi-family network hub across the order Bacteroidales (Figure 3c), serving as the primary reservoir for active Bacteroidales prophages (8/25 SA-vOTUs; Figure 5f) and co-clustering with the globally prevalent VFC_3 (Figure 4c). Collectively, these features identify SA-vFAM1 as a major, widely distributed and significantly activity-enriched lineage in GPAC.

While a minor subset of genomes exhibited weak similarity to Winoviridae, >96% of SA-vFAM1 members (4431/4583) lacked any ICTV-level classification (Figure S6e, Table S9). TerL-based phylogenetic reconstruction supported the evolutionary independence of SA-vFAM1 (Figure 5f). Whereas canonical Winoviridae references formed a restricted, peripheral clade, SA-vFAM1 members diversified into deep-branching lineages lacking classified representatives, indicating an expansive reservoir of uncharacterized viral diversity that lies beyond current ICTV representation. This topology mirrored our proteomic network boundaries, resolving discrete clades corresponding to subfamilies (e.g., vSUBFAM_4) and genera (e.g., vGENUS_8 and vGENUS_12). Notably, AvOTUs were distributed across diverse sub-lineages with frequent cross-genus host distribution. Among them, 8 SA-vOTUs were identified and concentrated in vSUBFAM_4 and vSUBFAM_26 (Figure 5g). Comparative genomics revealed that sequence conservation across these active lineages was primarily anchored by head–neck structural modules and conserved MuA/B transposition operons (Figure 5g). Diversity-generating retroelement (DGR) cassettes and three nucleotide metabolism/modification genes were also conserved across these subfamilies (Figure 5g). DGRs have been reported to diversify tail-fiber genes and thereby modulate host range in other systems^[16, 56–59]^. Their pervasive presence in SA-vFAM1 may point to a role in shaping the broad, multi-family host associations observed here. Within this conservation, TerS-TerL-adaptor complexes from diverse subfamilies clustered into a single homologous protein group, indicating strong phylogenetic conservation of the core packaging mechanisms.

Intriguingly, this distinctive suite of genomic hallmarks echoes descriptions of the informally proposed "hankyphages"^[60, 61]^. Querying the reference genome p00 via conserved TerL supported this link^[61]^, identifying 1,542 hanky-like phages and resolving their taxonomic placement, with 99.3% (1,531/1,542) mapping specifically into vSUBFAM_4 (Figures 5f, g, and S6e). Phenotypically, hanky-like phages exhibited a markedly higher active fraction compared to the family baseline (Figure 5e), a feature consistent with vSUBFAM_4 (10.5% active, 260/2,468). Furthermore, hanky-like prophage activity displayed pronounced Bacteroides specificity (Figure 5e). These patterns further support the presence of lineage-structured activity across both host and viral lineages. Moreover, the consistent activity of hanky-like phages across independent culturomics datasets further supports their broad activation potential across host contexts (Figure S6f).

In summary, SA-vFAM1 represents a hankyphage-inclusive prophage family associated with gut Bacteroidales. We propose the candidate name *Candidatus Mubacviridae* for this viral family, defined by: 1) a putative transposable lifestyle driven by a conserved MuA/B integration module; 2) conserved DGR cassettes potentially associated with broad host range; and 3) a structurally conserved head–neck and packaging module.

## Discussion

Prophages were detected in ∼80% of the cultured gut bacterial isolates analyzed here, yet their activity has remained difficult to assess. ACTIVE addresses a key limitation of existing coverage-based methods by using genome-wide host backgrounds rather than requiring flanking host sequence on the prophage-containing contig. It further combines nonparametric depth comparisons with a stringent effect-size threshold, reducing distributional assumptions while supporting conservative activity calls^[35]^. Together, these features extend activity assessment to fragmented isolate assemblies across large isolate sequencing collections. Beyond gut prophages, this framework would be applicable to prophages across diverse bacterial taxa and other replicative mobile genetic elements^[34, 62]^, providing replication evidence to prioritize candidates for downstream assays of progeny production and infectious particle release.

Distinct from sequence-only virome repositories^[14–17]^, the resulting Gut Prophage Activity Catalog (GPAC) links prophage sequences to cultured bacterial hosts and replication-associated activity annotations. By linking cultured bacterial hosts to metagenome-derived gut viruses, GPAC provides experimentally tractable entry points into previously uncharacterized viral diversity. This is well exemplified by the Bacteroidales-associated family SA-vFAM9 and three novel Mu-like families. Moreover, mapping activity profiles onto viral phylogenies pinpointed lineages markedly enriched for active members. Importantly, several of these active lineages align with globally prevalent gut viral clusters^[50]^, exemplified by the *Ca. Heliusvirales*-associated SA-vFAM7 and the Bacteroidales-associated Mu-like family SA-vFAM1 (*Ca. Mubacviridae*). Finally, GPAC provides isolate-resolved host associations, enabling systematic quantification of the observed host breadth and showing that active families such as SA-vFAM1 span multiple host families within Bacteroidales.

Our dataset reveals distinct prophage activity patterns structured by host taxonomy and viral lineage. Despite widespread lysogeny across gut bacteria, active prophage carriage was most strongly enriched in Bacillota rather than in Bacteroidota or Pseudomonadota, even though the latter phyla also harbored important active viral lineages. This pattern highlights extensive host–prophage interactions within Bacillota and prioritizes taxa such as Lactobacillaceae, Streptococcaceae and Enterococcaceae for dissecting how host intestinal physiology regulates the lysogeny–lysis decision^[63]^. Across the viral phylogeny, replication activity was concentrated within 46 SA-vFAMs with distinct intra-family architectures. Activity in SA-vFAM7 was restricted to a few SA-vOTUs, whereas SA-vFAM1 encompassed numerous phylogenetically diverse AvOTUs.

Moreover, SA-vOTUs more frequently exhibited cross-host distributions, particularly within SA-vFAM1, consistent with metagenomic observations of cross-family phage dynamics in the human gut. Extreme cases such as SA-vOTU4771, which showed putative cross-phylum distribution, echo sporadic metagenomic reports but require further experimental validation^[1, 64, 65]^.

Importantly, GPAC establishes a large-scale reference baseline for prophage replication activity under non-induced *in vitro* culture conditions. In GPAC, 2,920 prophages (5.4%) met our activity criteria across ∼10.5% of lysogens and ∼10.8% of vOTUs, generally with modest PtoH ratios (1.69-2.80×). This aligns with observations by Dahlman et al.^[21]^ in a smaller cultured cohort, where ∼4.9% (36/736) of gut prophages showed spontaneous induction under standard culture. It is also consistent with surveys by Wirbel et al.^[1]^, which reported that ∼5% of prophages displayed active dynamics in the gut with low-level induction (1-3× PtoH ratios). However, as reported by Dahlman et al.^[21]^ and Hu et al.^[66]^, prophages remaining dormant under standard conditions can be widely triggered by dietary compounds, epithelial secretions, or disease-associated metabolites, activating up to 22.63% of tested prophages and 36% of vOTUs. As additional gut-associated induction cues continue to be uncovered^[11, 67–69]^, These findings underscore that dormancy during standard culture does not imply a loss of inducible potential. Through a large-scale analysis that distinguishes lineages with elevated replication activity from those that remain dormant in vitro, GPAC provides prior information for designing matched induction experiments across host taxa and environmental cues to identify prophages with the strongest ecological impact.

In summary, by coupling an activity inference framework tailored for isolate sequencing with a high-resolution prophage atlas, this study systematically delineates the scale, evolutionary lineages, and host associations of active prophages across cultured gut bacteria. This activity-annotated resource will help guide future investigations into prophage induction dynamics and their broader ecological roles in the human gut.

## Limitations

Several limitations of this study warrant consideration. First, inherent to all coverage-based frameworks, ACTIVE captures DNA replication-associated enrichment rather than downstream biological endpoints; confirming active transcription, virion morphogenesis, and host lysis requires orthogonal validation^[61]^. Second, reliance on bulk sequencing depth precludes the detection of stochastic or ultra-low-frequency spontaneous induction events that fall below population-level resolution^[32]^. Third, while ACTIVE performs robustly on isolate genomes, its application to short-read metagenomes is subject to upstream assembly and binning artifacts^[70]^.

Thus, long-read or hybrid-assembled MAGs are strongly recommended for complex communities^[1]^.

## Methods

### Bacterial isolates

The *Acinetobacter baumannii* strains were previously isolated from sputum and cultured in this laboratory. The GBIC strains originated from a limited gut culture cohort previously isolated in the same laboratory. Genomic and sequencing data are available at PRJEB67733. All sequencing data underwent quality control using FastQC and Trimmomatic^[71]^ (SLIDINGWINDOW:4:25, MINLEN:100). Genomes were assembled using MEGAHIT^[72]^.

Five public gut culture cohorts with raw sequencing data and detailed metadata from PRJEB23845 (HBC)^[27]^, PRJNA482748 (CGR)^[28]^, PRJNA544527 (BIOML)^[26]^, PRJEB67733 (GBIC) and PRJNA1095184 (ATLAS)^[30]^ were utilized to evaluate gut prophage activity on a large scale. CheckM2^[73]^ evaluated genome quality. Fragmented genomes (longest contig length < 50 kbp, completeness < 40, contamination > 10) were excluded, yielding 31,150 genomes.

### Species tree of gut isolates

Taxonomic classification utilized GTDB-Tk^[74]^. Following a previously established two-step scalable species-level clustering framework^[75]^, all genomes were first partitioned by their assigned taxonomic orders using *classify_wf* modules of GTDB-Tk. Pairwise genomic distances within each order were rapidly estimated using Mash^[76]^ (-s 10000). Primary clusters were generated via average-linkage hierarchical clustering (UPGMA) using the fastcluster^[77]^ package with a distance threshold of 0.10 (corresponding to ∼90% ANI). For each primary cluster containing multiple genomes, pairwise precise ANI and alignment fractions (AF) were computed using skani^[78]^. Genomes sharing ≥95% ANI with ≥60% alignment coverage were defined as a single species cluster. Within each cluster, the genome with the highest quality score (S=Completeness−5× Contamination+0.5 × log_10_(N50), derived from CheckM2) was selected as the species representative. The taxonomic classification of the representative genome was assigned to all member genomes within the same cluster; unclassified lineages at any taxonomic rank were designated as novel taxa (e.g., ’new_species_1’). Finally, bacterial single-copy marker genes (bac120) were identified and aligned using the *identify* and *align* modules of GTDB-Tk. An approximate maximum-likelihood phylogenetic tree was inferred using FastTree (-WAG)^[79]^ with *Minisyncoccus archaeiphilus* (GCF_047159735) as the outgroup. Visualization used tvBOT^[80]^.

### Prophage induction

Bacterial strains were grown in LB broth at 37 °C with shaking (200 rpm) until the optical density at 600 nm (OD_600_) reached 0.5 (early exponential phase). Prophage induction was triggered by the addition of Mitomycin C (MMC) at a final concentration of 0.5 μg/mL (Sigma-Aldrich). An equal volume of the bacterial culture without MMC served as the control. Cultures were incubated for an additional 24 hours and then harvested.

### DNA extraction and sequencing

The bulk genome DNA was extracted from the cultures using the TIANamp Bacteria DNA Kit (Tiangen biochemical technology) following the manufacturer’s protocol with a minor modification: 200 μL of the culture was directly processed starting from the Proteinase K treatment step (Step 3). The genomic DNA was then sequenced on the Illumina NovaSeq 6000 platform using a 150-bp paired-end protocol.

For identification of induced prophages by virus-like particle sequencing (VLP-Seq), induced cultures were collected and centrifuged (5,000 rpm, 10 min) to remove bacterial cells. The supernatant was collected and filtered through a 0.22 μm filter. Viral particles were then precipitated with 0.5 M NaCl and 10% PEG 8000 at 4 °C overnight and resuspended in 500 μL of the corresponding culture medium. Subsequently, 2 μL of Thermo TURBO DNase and RNase were added, and the samples were incubated at 37 °C for 60 min, followed by enzyme inactivation at 95 °C for 15 min. Viral DNA was extracted using the TIANamp Virus DNA/RNA Kit (Tiangen Biotech) and sequenced on the Illumina NovaSeq 6000 platform using a 150 bp paired-end protocol.

Paired bulk and VLP samples were derived from the same 40 ml induction culture: 200 µL for bulk sequencing and the remaining supernatant for VLP enrichment, ensuring same-sample, same-batch pairing.

### Transmission electron microscopy (TEM)

Phage samples resuspended after PEG precipitation were negatively stained with 2% phosphotungstic acid (PTA). A 30 µL aliquot of the phage sample was applied to a copper grid and allowed to adsorb for 10 min. Excess sample was blotted off with filter paper, and 30 µL of 2% PTA staining solution was applied to the grid for 2 min. Excess stain was then blotted off, and the grid was dried under an infrared lamp for 10 min. Samples were observed using a JEOL-2100F transmission electron microscope operated at 120 kV.

### Prophage prediction

Seqkit^[81]^ extracted contigs > 2 kbp from all bacterial genomes. geNomad^[82]^, VIBRANT^[83]^, and VirSorter2^[84]^ predicted prophages. CheckV^[85]^ evaluated all predictions. Sequences with completeness ≥ 50 were retained and aligned to source genomes via blastn^[86]^. For each genome, blastn hits were ranked by score, and the highest-scoring hit matching the expected reference identifier was retained to determine exact coordinates. Sequences with overlapping intervals on the same reference contig were considered redundant (Figure S2a). For overlapping predictions from multiple tools, the sequence with the highest CheckV completeness was retained. The final deduplicated prophages and their coordinate files were used for genomic and activity analyses. Prophages located < 100 bp from either contig end were defined as terminal phages. All others were defined as internal phages. Terminal phages unanalyzable by PropagAtE were termed inaccessible terminal phages.

### Prophage activity

#### 1) Workflow of ACTIVE

ACTIVE takes the host genome, predicted prophage coordinates, and sequencing data as inputs. Read depths are extracted by mapping sequencing reads to prophage and host reference regions, yielding comparable depth distributions. ACTIVE operates in two modes: 1) *Precise* mode: Uses BEDTools^[87]^ to extract all host genomic regions outside the prophage as the background (non-phage in Figure 1a). 2) *Fast* mode: Uses host conserved single-copy marker genes (identified by Prodigal^[88]^ and fetchMGs^[38]^, default score > 300) as the background (MGs in Figure 1a). ACTIVE first assesses the normality of depth distributions to select statistical tests (Figure S1b). A one-sided upper-tail non-parametric test was used to determine whether depth in prophage regions was significantly greater than that in host reference regions, with effect size used to quantify the magnitude of the difference. Parametric tests were performed only when both prophage and host depth distributions passed the normality assessment. The vast majority of sequencing datasets were better suited to the more conservative non-parametric test. All analyses in this study also applied the non-parametric Brunner–Munzel test. Prophages required a minimum mapping coverage ratio (default 0.75). For significant prophages (p-value < 0.01) in the upper-tail test, ACTIVE calculates the Cliff’s Delta as effect size. To minimize false positives, active prophages were defined as having a p-value < 0.01 and Cliff’s Delta ≥ 0.8. To prevent R integer overflow from large host genomes (2-5 Mbp) when calculating Cliff’s Delta in *Precise* mode, ACTIVE approximates the full-genome effect size using Monte-Carlo subsampling (Figure S1c). In each of 500 iterations, a subset of 10,000 individual host depth positions is drawn without replacement from the host genomic background, and Cliff’s Delta is calculated between the full prophage depth vector and this host subsample. The mean across the 500 iterations is used as the estimated full-genome Cliff’s Delta and is the only quantity compared against the activity threshold (δ ≥ 0.8) (Figure S1d). Read mapping utilized BBMap^[89]^ (ambiguous = random, minid = 0.95) for short reads and minimap2^[90]^ for long-read sequencing. SAMtools^[91]^ and BEDTools extracted depth data.

#### 2) PropagAtE, Prophage Tracer and mVIRs

Bulk sequencing data were processed using the default pipelines. For PropagAtE^[35]^, prophages lacking a host reference background (host_len=0) for comparison were defined as inaccessible terminal phages. By default, prophages required a minimum mapping coverage ratio = 0.5. PropagAtE classified a prophage as active when Cohen’s D ≥ 0.7 and prophage-to-host ratio (PtoH) on prophage-containing contig was ≥ 2. For Prophage Tracer^[32]^, we defined two custom modes based on SR/DRP evidence counts. In full mode, a prophage was classified as active if any SR/DRP evidence was detected. In strict mode, active classification additionally required at least one SR or DRP evidence count ≥20. For mVIRs^[31]^, the filtered FASTA files output by the software were considered active prophages.

### Benchmarking on complete bacterial genomes (Benchmark 1)

To evaluate the performance of ACTIVE against existing computational tools, we curated a comprehensive ground-truth reference dataset comprising 348 experimentally validated prophage induction events across diverse bacterial species. Each event was defined by: (1) a complete, circularized host bacterial reference genome; (2) curated genomic coordinates of integrated prophages; and (3) publicly available bulk whole-genome sequencing (WGS) data generated under distinct induction regimens (Table S1). All host genomes, prophage coordinates, sequencing datasets, and prophage activity states (used as ground truth) were collected from the corresponding published literature (Table S1). For each validation event, raw sequencing reads were quality-trimmed using Trimmomatic (SLIDINGWINDOW:4:25 MINLEN:100). All events were then predicted with ACTIVE, PropagAtE, Prophage Tracer, and mVIRs using their default parameters. The BAM file produced by ACTIVE in *Precise* mode was reused for both the *Fas*t mode and PropagAtE. Because Prophage Tracer and mVIRs output coordinates rather than per-prophage activity labels, a predicted prophage was counted as a true positive when it overlapped a ground-truth prophage on the same host contig by ≥50% of its own length.

To ensure a fair comparison, benchmarking was restricted to the intersection of evaluable events across tools, defined as the Callable Intersection: prophages that passed both the ACTIVE and PropagAtE coverage thresholds and satisfied PropagAtE’s requirement of host_len > 0. Events failing these criteria were labelled QC_FAIL.

For each tool or mode, TP, FP, TN, and FN were counted on the Callable Intersection, and sensitivity, specificity, precision, accuracy, F1, and Matthews Correlation Coefficient (MCC) were derived accordingly. To assess robustness to the ACTIVE decision threshold, the significance label was held fixed while the Cliff’s Delta threshold was varied from 0 to 1 in steps of 0.001, and sensitivity, specificity, precision, F1, MCC, and accuracy were recomputed at each threshold. Receiver operating characteristic (ROC) and precision–recall (PR) curves were generated from the continuous Cliff’s Delta scores, and the area under the ROC curve (AUC) and the area under the PR curve (AUC-PR) were calculated.

### Benchmarking on fragmented bacterial genomes (Benchmark 2)

The complete bacterial reference genomes in Benchmark 1 were originally derived from hybrid assemblies combining long-and short-read sequencing. To evaluate tool performance on fragmented draft assemblies typical of routine isolate sequencing, we constructed a matched, isogenic short-read draft counterpart for each Benchmark 1 isolate. Raw Illumina reads retrieved from the corresponding isolates were quality-trimmed using Trimmomatic (SLIDINGWINDOW:4:25 MINLEN:100) and assembled *de novo* using MEGAHIT.

Verified prophage sequences from complete genomes were mapped onto draft contigs using BLASTN (-task megablast -evalue 1e-10 -perc_identity 95.0), and alignments on the same contig separated by ≤1,000 bp were merged. On the basis of strict alignments (≥99% identity; designated p99), prophages were hierarchically classified as split (≥2 contigs), gapped (disjoint intervals on a single contig), intact (single contig with ≥90% query coverage), or partial (query coverage <90%) (Figure S1f). High-confidence assembled prophages with cumulative p99 query coverage ≥80% were compiled into the final evaluation set, and the prevalence of prophages located at contig boundaries (terminal prophages) within this set was quantified.

The activity of prophages in this evaluation set was predicted using ACTIVE under both *Precise* and *Fast* modes and evaluated against the ground truth to derive standard classification metrics on fragmented genomes. For prophages split across multiple contigs, per-base depths across all mapped segments were concatenated into a single depth vector for subsequent comparison against the host genomic background. To compare the effective computational capacity (callable coverage rate) between ACTIVE and PropagAtE, both tools excluded prophages with a breadth of coverage ≤ 0.75; for PropagAtE, prophages on contigs lacking flanking host sequences (host_len = 0) were additionally excluded.

### Descriptive statistics of host prophage-associated traits

We characterized host prophage-associated traits using three core parameters: lysogeny prevalence, active prophage carriage prevalence, polylysogeny prevalence. Two calculation methods were employed.

1) Unweighted Rate: Calculated as the pooled average across all isolates.

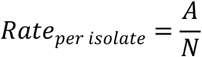

Lysogeny prevalence was the proportion of isolates carrying at least one prophage. Active prophage carriage prevalence and polylysogeny prevalence were the proportions of lysogenic isolates carrying at least one active prophage and more than one prophage, respectively.

2) Taxonomic Weighted Rate^[92]^: Calculated as the arithmetic mean of rates across K distinct taxa (e.g., species) to eliminate bias from species overrepresentation:

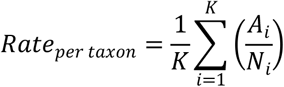

For each taxonomic unit *i*, A_i_ represents the number of positive events (genomes with prophage for lysogeny prevalence, genomes with > 1 prophages for polylysogeny prevalence, or genomes carrying active prophages for active prophage carriage, respectively), while N_i_ denotes the corresponding denominator population (total genomes for lysogeny prevalence; total lysogens for polylysogeny prevalence and active prophage carriage). Population weighted averages summarized these rates per phylum.

### Hierarchical generalized linear mixed models for host prophage-associated traits

We used the R package lme4 to model three isolate-level traits. Prophage carriage was analyzed across all isolates using a binomial generalized linear mixed model (GLMM) with a logit link. Among lysogenic isolates, active prophage carriage was analyzed using a second binomial GLMM, whereas natural-log-transformed prophage counts were analyzed using a Gaussian linear mixed model. All models included standardized genome completeness and log_10_-transformed contig N50 as fixed covariates, with random intercepts for isolation source and nested host taxonomy: (1 | Phylum/Class/Order/Family/Genus/Species). Variance components quantified variation associated with source and taxonomic levels. Component proportions were normalized to total random-effect variance for binomial models and additionally included residual variance for the Gaussian model. Conditional random-intercept estimates and their conditional standard deviations summarized rank-specific taxonomic deviations. Ranking plots included taxa represented by at least five isolates in the corresponding analysis subset.

### Genomic and phenotypic divergence of host prophage-associated traits

For each species represented by at least five isolates and three lysogens, we calculated lysogeny prevalence and active prophage carriage prevalence. The latter was calculated among lysogens only; species without lysogens were excluded rather than assigned an active-carriage prevalence of zero. Phenotypic distance between congeneric species was the Euclidean distance between these two unscaled prevalences. Mean genomic distance was calculated from skani ANI across genome pairs. Their association was assessed using Spearman correlation, with significance evaluated by permutation to account for non-independent species pairs.

### Phage genome classification and clustering

1) Taxonomy assignment

The UHGV-classifier^[16]^ assigned UHGV clusters and ICTV taxonomy to all phages.

2) Clustering into vOTU based on genomic similarity

vOTU and higher-ranking clusters were established following previous methods^[16]^. An all-versus-all blastn search utilizing the megablast task processed all genomes. A CheckV script (https://bitbucket.org/berkeleylab/checkv/src/master/scripts/anicalc.py) calculated average nucleotide identity (ANI) and alignment fraction (AF). Following MIUViG guidelines^[93]^, genome connections required at least 95% ANI over 85% of the shorter sequence. Intergenome similarities were calculated as (ANI × AF) / 100. Clustering utilized an iterative tiered approach integrated with the Markov Cluster Algorithm (MCL). Sequences were categorized into three tiers based on CheckV assessments: complete genomes, high quality genomes (completeness ≥ 90), and fragments. MCL clustering (inflation 2.0) on the first tier established core vOTUs. Subsequent tiers were iteratively incorporated into existing or novel clusters using the identical ANI and AF thresholds.A hierarchical decision tree selected a single representative sequence for each vOTU cluster. CheckV verified complete genomes longer than the cluster median length received priority. For clusters lacking complete genomes, the representative minimized the difference between observed contig length and predicted total genome length while prioritizing higher viral gene counts.

3) Cluster into higher hierarchy with proteomic similarity

Representative genomes of species-level vOTUs were grouped into a taxonomic hierarchy based on whole-genome proteomic similarity. geNomad predicted open reading frames for all vOTU representatives.

DIAMOND^[94]^ executed an all-versus-all protein similarity search (very sensitive mode, e-value 0.001). Intergenomic similarity utilized a normalized bitscore metric, calculated as the sum of best reciprocal hit bitscores divided by the lower of the two self score totals. MCL defined hierarchical viral clusters across four increasing stringency thresholds: vFAM (0.055), vSUBFAM (0.32), vGENUS (0.65), and vSUBGEN (0.80). A top-down pruning strategy ensured taxonomic coherence. A similarity link at a higher stringency rank was retained only if both genomes shared the same cluster at the preceding lower stringency rank. Singletons received unique identifiers, resulting in a stable nested taxonomic hierarchy.

4) Co-clustering of vFAMs with reference Viral Family Clusters (VFCs)

Representative genomes from GPAC vOTUs and 12 previously characterized high-prevalence viral family clusters (VFCs; present in >50% of global gut viromes) were cross-clustered. ORFs for the reference VFC genomes were predicted using geNomad. Family-level clustering was then performed on the combined set of representative genomes as described above, generating a set of cross-dataset family clusters designated as NvFAMs (new viral family clusters). To assess the correspondence between each vFAM and reference VFC, we computed two complementary proportions: first, the fraction of vFAM member genomes co-clustering into the corresponding NvFAM; and second, the fraction of reference VFC member genomes captured within that NvFAM.

### Identification of significantly activity-enriched vOTUs (SA-vOTUs) and vFAMs (SA-vFAMS)

Active vOTUs (AvOTUs) and active vFAMs (AvFAMs) were defined as taxa containing at least one active genome. To identify lineages with exceptionally high activation frequencies, statistical analyses were performed at the AvOTU and AvFAM levels using a source-stratified exact framework (minimum 5 genomes per taxon). For taxa distributed across multiple sources (i.e., different culture origins), the Cochran–Mantel–Haenszel (CMH) test was applied, controlling for “source” as a stratifying variable. For taxa detected in only one source, Fisher’s exact test was used against the local background baseline. Effect size was quantified using the Mantel–Haenszel common odds ratio (OR) with Haldane–Anscombe correction. Taxa were designated as significantly activity-enriched units (SA-vOTUs and SA-vFAMs) only if they met: global FDR (Benjamini–Hochberg) < 0.05 and common OR > 1.

Normalized rarefaction curves assessed sampling saturation for total vOTUs, SA-vOTUs and SA-vFAMs across host phyla. A random subsampling procedure (10 iterations per step, 30 steps total) evaluated each host phylum containing at least five genomes.To facilitate cross-phylum comparison, the sampled host genome count and the cumulative viral entity count were normalized. LOESS smoothing fitted the resulting curves.

### Cross-Host distribution and host-virus interaction networks

The host distribution breadth for each vOTU was defined by the highest taxonomic rank at which divergence occurred among its source hosts, ranging from Cross-Species to Cross-Phylum. A vOTU was defined as cross-host if its host distribution spanned at least the species level (i.e., cross-species or higher).

Bipartite interaction networks connecting hosts and AvOTUs were constructed. The taxonomic breadth of host distribution categorized the viral host distribution. To display viral connectivity across host reservoirs, networks of shared vOTUs, regardless of activity status, were constructed at the family, genus, and species levels.

Connections between host nodes were established if they shared vOTUs. Edge weights represented the number of shared vOTUs. The proportions of cross-host vOTUs among inactive vOTUs, SA-vOTUs, and other AvOTUs (not significant or excluded AvOTUs described above) were compared using Fisher’s exact test with FDR correction.

### Genetic distance of prophage

To identify the closest reference genome for each prophage, pair-wise genetic distances were computed using BinDash2^[95]^ against established viral databases, including INPHARED^[46]^ (7Apr2026 version), ProphageDB^[18]^, and the Unified Human Gastrointestinal Virome (UHGV)^[16]^. For prophages belonging to putative Mu-like vFAMs, comparisons were specifically performed against the Transposable Bacteriophage Genome Database (TBPGD)^[55]^.

### Prophage phylogenetic trees and genomic comparisons

For TerL-based phylogenetic tree of all *Caudoviricetes* vOTU in GPAC, to reliably represent each vOTU in phylogenetic analyses, the terminase large subunit (TerL) amino acid sequences were retrieved, excluding genomes classified as low quality. For each vOTU, a single-copy TerL gene from the cluster representative was prioritized. If the representative genome lacked a TerL gene or harbored multiple copies, a valid constituent member genome encoding exactly one TerL gene was selected as a proxy. vOTUs lacking an unambiguous single-copy TerL across all members were excluded from phylogenetic inference. vOTU TerL sequences were merged with reference sequences from the ICTV *Caudoviricetes.* For TerL-based phylogenetic tree of SA-vFAM1 vOTU TerL sequences were extracted as described above. Instead of including ICTV Caudoviricetes, the Hankyphage reference strain p00 (BK010646) and three *Winoviridae* (MT732475, MN812203, MN812205) were used as the reference^[61]^, with *Carjivirus communis* (BK010471) from *Crassvirales* as the outgroup. For TerL-based phylogenetic tree of vFAM7, the Pharokka annotations facilitated the extraction of TerL protein sequences from the genomes, excluding those containing multiple

TerL copies. All TerL-based phylogenetic trees were constructed by aligning TerL sequences with MAFFT^[96]^, followed by phylogenetic reconstruction using FastTree with the WAG model (-wag).

For vFAM9, ViPTree^[97]^ constructed the whole proteome phylogenetic trees with default reference.

Clinker^[98]^ executed the protein-based comparative genomic analyses. vConTACT2^[99]^ constructed the proteome similarity network.

### Prophage annotation

Pharokka^[47]^ was used to annotate and plotted prophage genomes. Three essential morphogenetic genes, the TerL, portal protein, and major head protein, were profiled across Caudoviricetes genomes and clustered vOTUs. A vOTU was defined as positive if the marker was detected in ≥1 constituent genome.

For vFAM7, four core components constituting the packaging module were further analyzed: HNH endonuclease, TerS, TerL, and portal protein. To ensure structural completeness and proximity, a distance-based search identified valid modules with an HNH-TerS interval of 17 genes or fewer and a portal protein within five genes of TerL. For genomes with multiple candidates, the most complete and compact module was prioritized as the representative locus. Functional annotations of genes within these defined intervals were extracted, excluding non-informative terms such as "hypothetical protein" or "unknown function". Finally, frequency analysis quantified the occurrence of specific protein domains and functional categories.

### Identification of enriched genetic elements within active prophages

Enriched genetic elements within active prophages were identified using Pharokka annotations and protein sequences from high-quality (CheckV completeness ≥ 90) *Caudoviricetes* genomes. This analysis comprised both term-based and domain-based enrichment.

For term-based enrichment, a two-step statistical modeling approach was employed to identify features significantly associated with prophage activity. Terms present in fewer than 1% of the total modeled prophages, as well as those annotated as hypothetical proteins or unknown functions, were excluded. First, a binomial generalized linear model (GLM) screened for terms associated with prophage activity (FDR < 0.05), adjusting for host phylum, prophage location type, and genome length as fixed effects. Next, a generalized linear mixed model (GLMM) validated these candidates by incorporating the same fixed effects alongside vFAMs as a random effect to control for phylogenetic clustering. Validated features (GLMM FDR < 0.05) were subsequently categorized as induced-enriched (log2 estimate > 0) or dormant-enriched (log2 estimate < 0).

For domain-based validation, protein sequences were clustered into protein clusters (PCs) using MMseqs2^[100]^ (--min-seq-id 0.3 -c 0.7 --cov-mode 0). Representative sequences of each PC were annotated via MetaCerberus^[101]^ against COG, PFAM, KOFam_prokaryote, PHROG, and VOG databases, with functional assignments subsequently extended to all cluster members. For each functionally annotated domain, we calculated its genomic proportion within each vOTU, defined as the number of genomes in the vOTU containing that domain divided by the total number of genomes assigned to the vOTU. To reduce stochastic noise, domains that exhibited a genomic proportion ≥ 40% in at least five distinct vOTUs were retained. Candidate domains were initially screened using quasi-binomial GLM, with the proportion of active genomes per vOTU as the response while the genomic fraction of the domain, host phylum, prophage location type, and vOTU size were included as predictors. Those with FDR ≤ 0.05 were then validated using a binomial GLMM that additionally included viral family as a random effect. Domains achieving an FDR < 0.05 were designated as strictly validated core domains. For these core domains, we compared the average active proportion between vOTUs with and without the domain.

### Identification of gut Mu-like transposable prophages

This analysis used high quality *Caudoviricetes* genomes (CheckV completeness ≥ 90%), as above. To identify integrases and transposases missed by Pharokka, an HMM search^[102]^ (-E 1e-5) was performed using specific Pfam profiles. Integrases were identified via PF07508, PF00239, and PF00589, while transposases were identified via PF00665, PF13333, and PF13683^[1]^. High-confidence transposase elements were further annotated using ISEScan^[103]^ (E ≤ 1e-09, score ≥ 60) to call those still unknown. CDS immediately flanking transposases (±1 CDS) with the same transcriptional orientation were examined and classified by PHROG and functional category.

Mu-like phages were specifically identified by their conserved MuA/B transposition modules, which function in place of standard integrases. MuB was annotated by searching PC representative sequences against the COG2842 database using rpsblast^[104]^ (-E 1e-5). A phage was classified as Mu-like if a transposase gene was located immediately adjacent to and in the same orientation as a MuB gene. To confirm that transposition was evolutionarily conserved rather than sporadic insertion, the prevalence of Mu-like modules was assessed across each vFAM. A vFAM was defined as Mu-like if at least 80% of its transposase-encoding members harbored the canonical adjacent MuA/B operon. The phylogenetic homology of candidate vFAM members was checked if there were hits with known Mu-like viruses in ICTV taxonomy using UHGV-classifier, such as the genus *Muvirus*.

### Identification of Hanky-like phage in SA-vFAM1

Following established methods^[61]^, SA-vFAM1 sequences were queried against the hankyphage reference genome p00 using blastn (-evalue 1e-5 -max_hsps 1). Candidate hankyphage-like prophages were subsequently identified from hits maintaining a minimum alignment coverage of 10%.

## Data analysis

Statistical analysis and plotting were performed using R 4.3.3.

## Data availability

Raw sequencing data generated in this study have been deposited in PRJCA063323. All source data, which include prophage raw activity results, bacterial taxonomy and all raw data of raw figures are available at FigShare.

## Code availability

All scripts for analysis and datasets needed to recreate all tables and figures in the manuscript, and ACTIVE script, are available at https://github.com/SIAT-MaLab/ACTIVE.

## Supporting information

Suppplementary figures

## Acknowledgments

We are grateful to Dr. Zepeng Qu at the Chinese Academy of Sciences for kindly providing GBIC gut strains collection. This research was supported by the National Key R&D Program of China (2024YFA0919400), the Shenzhen Medical Research Funds (B2302023), the National Natural Science Foundation of China (32301224), and the Guangdong Natural Science Foundation General Program (2024A1515012119).

## Author contributions

Conceptualization, Y.M., Y.H.; investigation, Y.H., Z.L., J.L.; resources, X.T., L.D.; writing—original draft, Y.H.; writing—review & editing, Z.L., J.L, J.H., X.T., L.D., and Y.M.; supervision, Y.M.

## Competing Interests

X.T. is CEO of Synphage Biotechnology. Y.M. is co-founder of Synphage Biotechnology. The other authors declare no competing interests. Some schematic diagrams were created with BioRender.com. Publication license obtained.

## Notes

### Summary of Updates

Add a benchmark and revise the manuscript without affecting the conclusions.

