## Supplementary material for "A genome-resolved atlas reveals lineage-specific activity patterns of prophages in cultured human gut bacteria": Suppplementary figures

Yingfei Ma

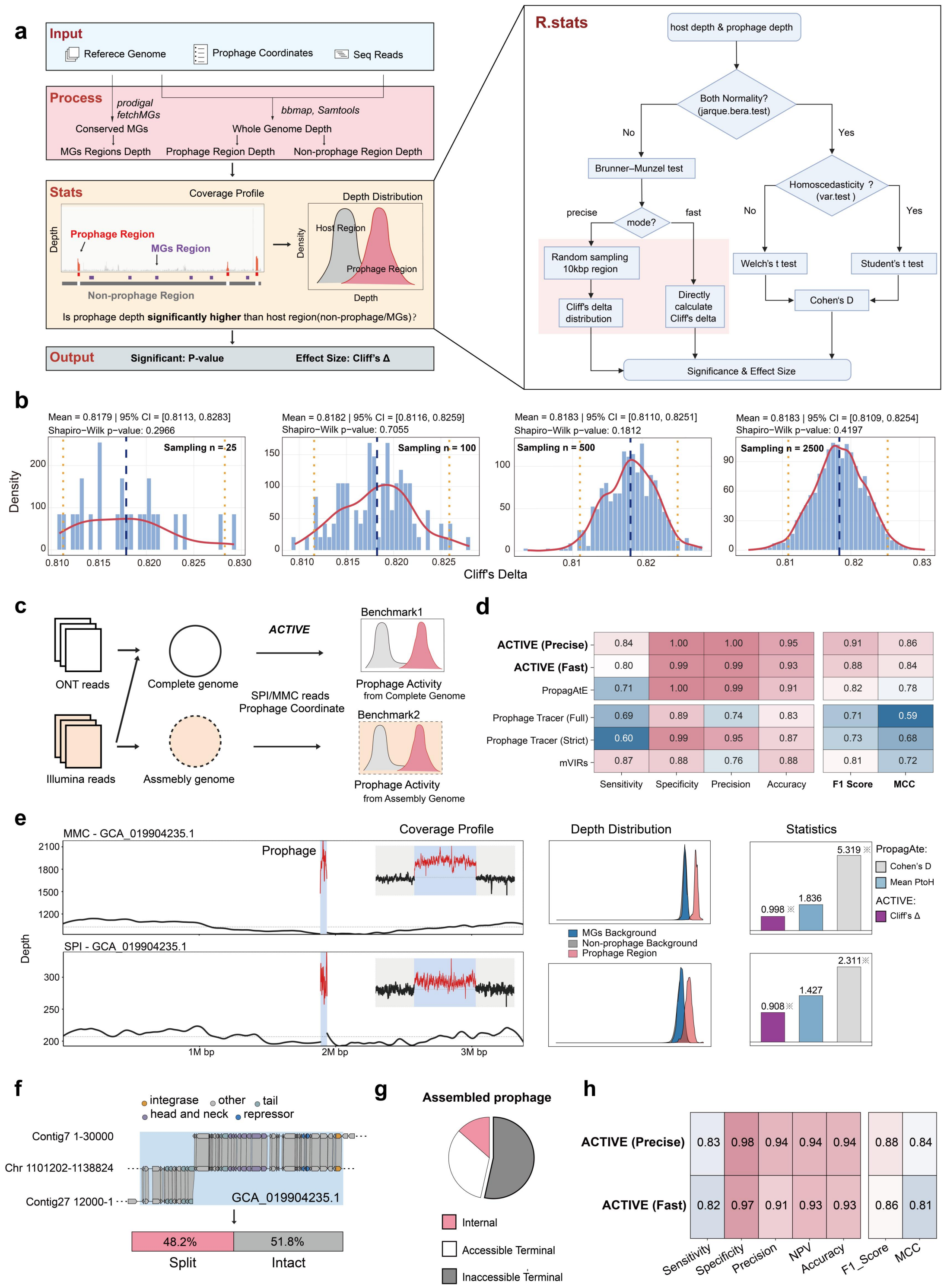


Figure S1. Workflow of ACTIVE and benchmarking performance across complete and fragmented genomes

(a) Workflow schematic. ACTIVE maps reads to host genomes to extract coverage distributions across prophages, non-prophage regions, and conserved marker genes (MGs). Significance and effect sizes are assessed by default using the Brunner–Munzel test and Cliff's Delta, with parametric alternatives supported (R.stats). Fast mode calculates Cliff's Delta directly, whereas Precise mode uses random subsampling across 10-kb host windows.

(b) Cliff's Delta distributions across repeated host subsampling (n = 25–2,500 iterations). Blue dashed lines: means (estimated full-genome effect size). Orange dotted lines: 2.5th–97.5th percentiles of the subsample values.

(c) Benchmark design. Benchmark 1 assesses predictions against experimental ground truth using complete hybrid-assembled genomes. Benchmark 2 evaluates the same loci across fragmented *de novo* short-read assemblies (Methods).

(d) Performance on complete genomes (Benchmark 1). Classification metrics of ACTIVE (Precise/Fast; Cliff's Delta ≥ 0.8) compared with PropagAtE, Prophage Tracer, and mVIRs.

(e) Representative discordant classification. Validated active prophage (GCA_019904235.1) correctly identified by ACTIVE (Cliff's Delta ≥ 0.8) but classified as dormant by PropagAtE due to baseline thresholds (PtoH < 2.0 or Cohen's D < 0.7). Panels show coverage profiles, depth distributions, and statistical outputs. Asterisks (*) indicate criteria met.

(f) Prophage fragmentation in Benchmark 2. Schematic and proportions of intact prophages (single contig; 51.8%) versus split prophages spanning multiple contigs (48.2%).

(g) Distribution of internal, accessible terminal (host flank > 0 bp), and inaccessible terminal prophages (host flank = 0 bp; lacking flanking sequences required by PropagAtE).

(h) Performance metrics of ACTIVE in Fast and Precise modes on Benchmark 2.

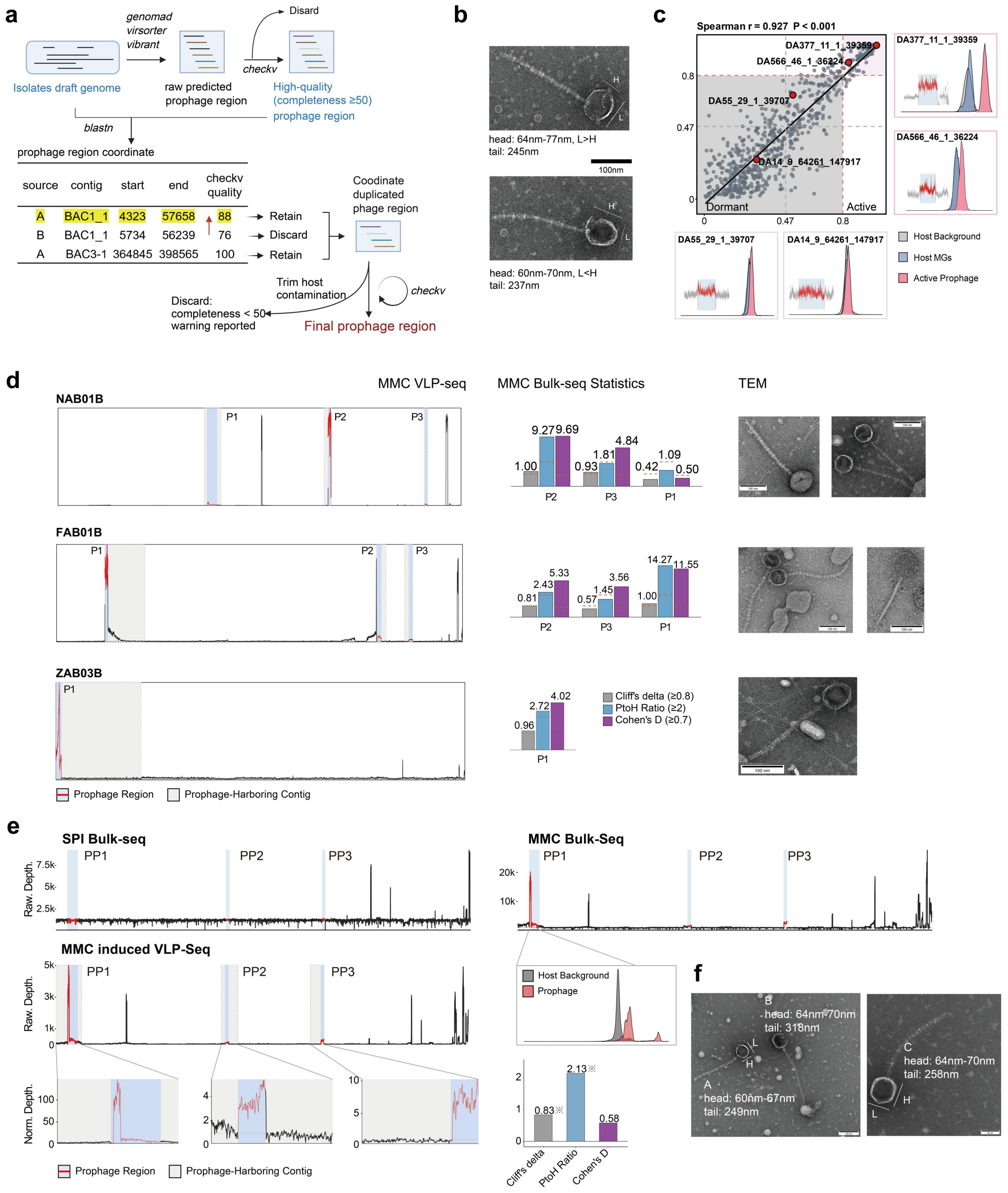


**g**

| prophage_candidate | contig | attL_start | attL_end | attR_start | attR_end | prophage_ | SR_evidence_attB | SR_evidence_attP | DRP_evidence_attB | DRP_evidence_attP |
|---|---|---|---|---|---|---|---|---|---|---|
| candidate_39 | SAB43B_1 | 114124 | 114133 | 128040 | 128049 | 13916 | 0 | 1 | 12 | 153 |
| candidate_62 | SAB43B_1 | 114049 | 114092 | 128173 | 128217 | 14124 | 0 | 3768 | 0 | 2132 |
| candidate_84 | SAB43B_1 | 128166 | 128197 | 210162 | 210193 | 81996 | 8 | 46 | 2 | 20 |

Figure S2. Prophage activity evaluation in the *A. baumannii* (AB) and GBIC datasets.

(a) Prophage identification and curation pipeline. Candidate prophage regions predicted by geNomad, VIBRANT, and VirSorter2 were deduplicated based on genomic coordinates. For overlapping regions, the prediction with the highest CheckV quality score was retained, followed by host contamination trimming and completeness filtering ( ≥ 50%; Methods).

(b) Transmission electron micrographs of virions in the MMC-induced virus-containing supernatant of strain RAB20B. Scale bar, 100 nm. The two examples show tailed phages with similar tail lengths but different head dimensions (L and H).

(c) Concordance between the Fast and Precise modes of ACTIVE across AB and GBIC isolates. Each point represents a prophage, and the scatter plot compares Cliff's Delta obtained with the two modes. Representative prophages in the Dormant and Active ranges are labeled. Corresponding depth distributions and coverage profiles of these representative prophages are shown. In the depth distributions, gray, blue and red denote the whole-genome host as background used in Precise mode, the host MGs as background used in Fast mode, and the prophage region, respectively. In the coverage profiles, gray indicates the flanking host region, whereas the red line and blue shading indicate the prophage region.

(d) Experimental validation in additional *A. baumannii* strains. Left: VLP-seq coverage profiles across predicted prophage regions under MMC induction. Middle: Statistical metrics computed from bulk-seq data using ACTIVE and PropagAtE; orange dashed lines indicate active thresholds. Right: TEM micrographs of virions released upon MMC induction for each corresponding strain.

(e) Experimental validation in strain SAB43B. Left and middle: genome-wide depth profiles across spontaneous induction (SPI bulk-seq), MMC induction (MMC bulk-seq), and MMC-induced virion-enriched fractions (MMC-induced VLP-seq), with blue shading indicating three predicted prophages (PP1–PP3). Bottom insets highlight prophage contigs (gray) and prophage regions (blue), shown as raw depth (bulk-seq) or median-normalized depth (VLP-seq). Right: read-depth density of PP1 relative to host background (top) and statistical metrics (asterisks indicate values meeting active thresholds). PropagAtE classifies a prophage as active when Cohen's D≥ 0.7 and PtoH ≥ 2, whereas ACTIVE uses Cliff's Delta ≥ 0.8.

(f) Transmission electron micrographs of induced virions in strain RAB20B (scale bars: left, 100 nm; right, 50 nm). Three distinct morphotypes (A–C) are shown, differing in tail length (A vs. B) or capsid dimensions (A vs. C; length L, height H).

(g) Prophage Tracer (Strict) predictions resolving PP1 into a PICI-like element (PP1-1, corresponding to candidate_39/62) and an intact prophage (PP1-2, corresponding to candidate_84).

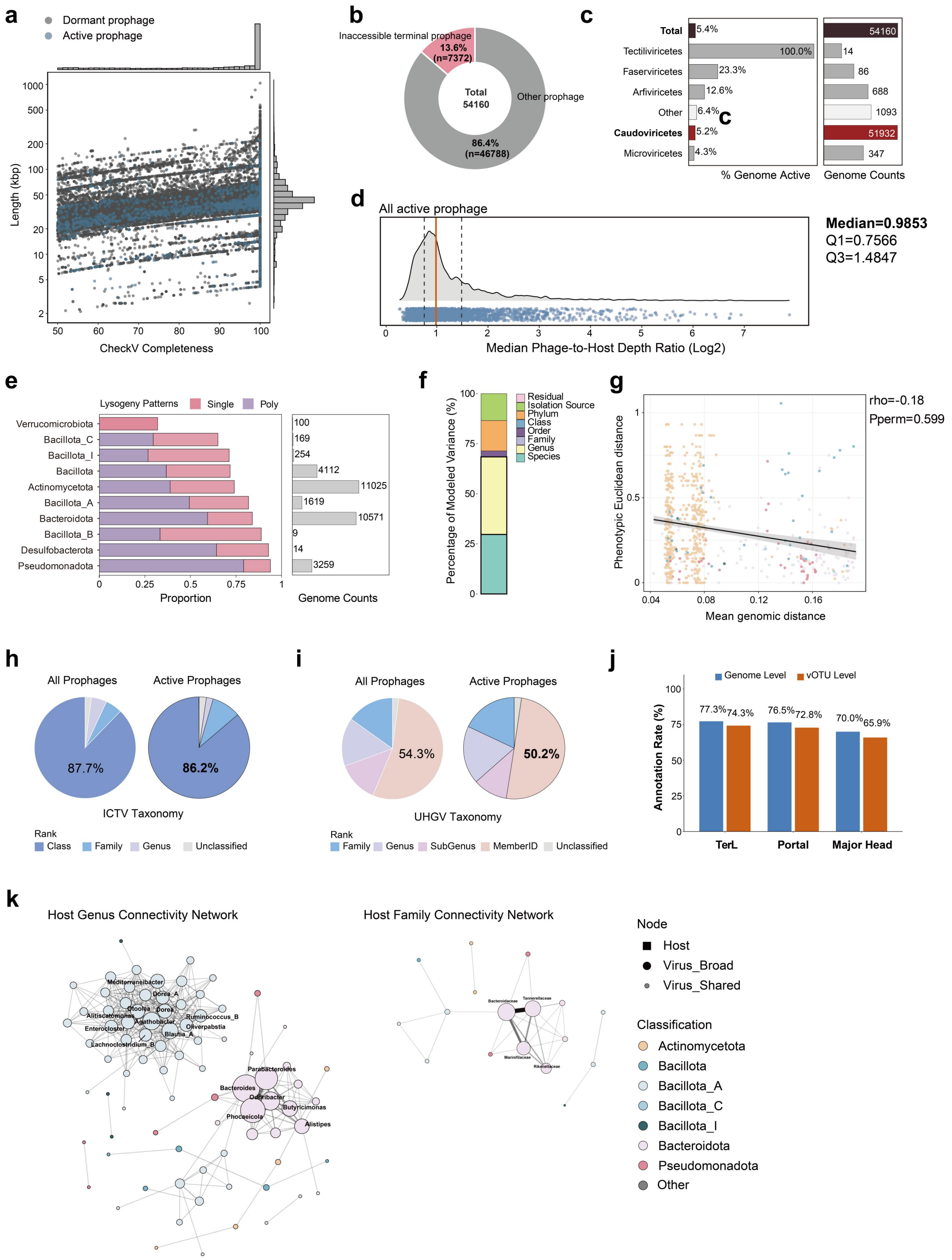


Figure S3. Profiling Prophages in the Gut Culturome.

(a) Prophage length and CheckV completeness. Distribution of active (blue) and dormant (gray) prophages. Marginal histograms show that lengths and CheckV completeness are concentrated at ~40 kbp and ~100% (complete), respectively.

(b) Terminal prophage configurations in ACTIVE-DB, highlighting prophages with host_len = 0, which are inaccessible to PropagAtE.

(c-d) Distribution of ICTV classes and their corresponding activation rate of all prophages (c), alongside median prophage-to-host ratio (mPtoH, median prophage region coverage/median host region coverage) of active prophages (d). Median: solid orange lines; quartiles: grey dashed lines. Median, Q1, and Q3 are log2-transformed values.

(e) Phylum-level summary of lysogeny of gut isolates, displaying the relative abundance of non-lysogens, mono-lysogens, and polylysogens in relation to the genomic sample size.

(f) Variance decomposition across phylogenetic hierarchies. The proportion of total phenotypic variance in active prophage carriage explained by each nested taxonomic level (from phylum down to species) is shown.

(g) Divergence of prophage-associated traits relative to genomic distance. Scatter plot showing the relationship between genomic distance (1 - ANI) and prophage-associated traits for intra-genus species pairs (Methods). Each point represents a pairwise comparison between two species within the same genus, colored by phylum. The regression line (black) with a 95% confidence interval (gray) illustrates the overall trend. Statistics for individual phyla are provided in the legend.

(h-i) The lowest assigned taxonomic units for all prophage genomes or active genomes according to both ICTV (h) and UHGV (i) classification systems.

(j) Proportion of *Caudoviricetes* prophages annotated with three conserved packaging and structural proteins (TerL, portal, and major head) using Pharokka. Rates are evaluated at the genome and vOTU levels; a vOTU is scored as annotated if ≥1 member genome is annotated.

(k) Genus-level and family-level host connectivity networks. Edges denote shared vOTUs between host taxa.

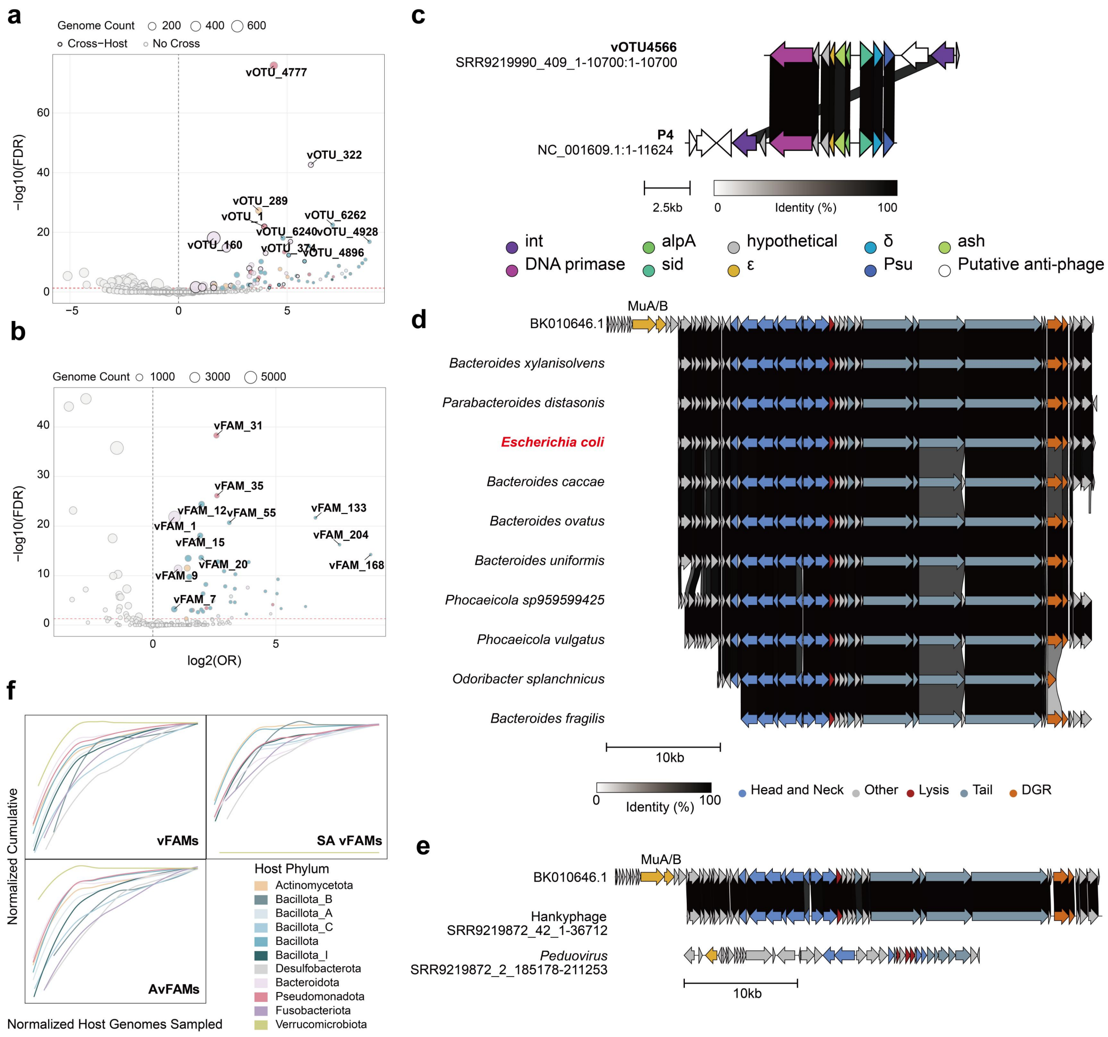


Figure S4. Identification of significant active prophage lineages.

(a,b) Volcano plots of activity landscape of AvOTU (a) and AvFAM (b) levels. x- axis: $\log_2$(odds ratio); y- axis: $-\log_{10}$(FDR). Significant activity-enriched taxa are highlighted by host phylum. Bubble size indicates genome count of viral taxon. Black borders indicate cross- host prophages. Dashed lines: FDR = 0.05 (red) and $\log_2$OR = 0 (black). Top 10 candidates by effect size and significance are labeled.

(c) vOTU4777, the most significant SA-vOTU, is a P4-like satellite phage.

(d-e) A Cross-Phylum SA-vOTU. (d) vOTU4771 members from different hosts genera, indicating that phage from *E.coli* is highly similar to others from Bacteroidales. (e) *E. coli* strain SRR92198672 harbors a hankyphage from vOTU4771 alongside a genomically distinct *Peduovirus*.

(f) Accumulation curves of vFAMs, AvFAMs, and SA-vFAMs across host phyla.

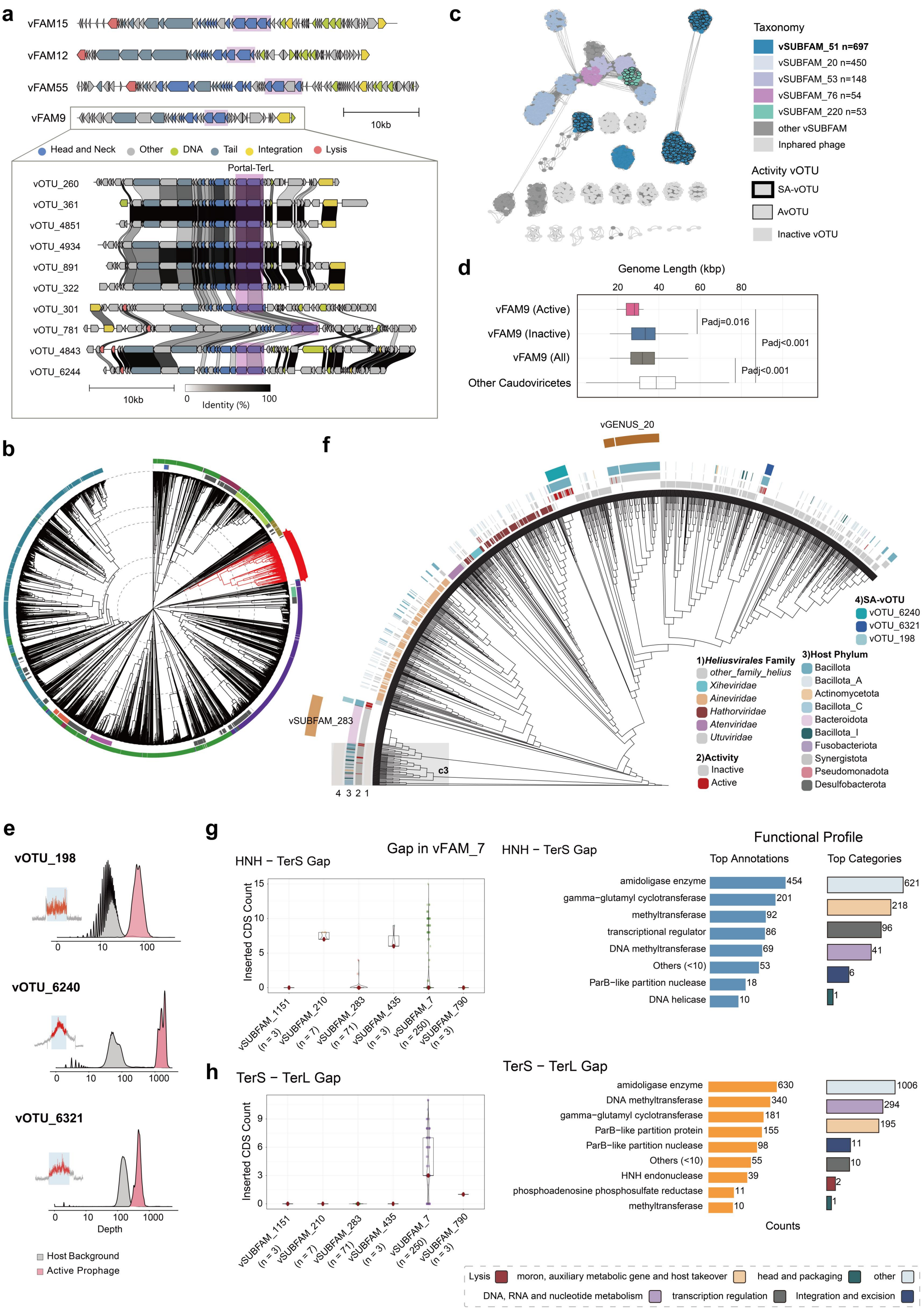


Figure S5. vFAM9 and vFAM7 are SA-vFAMs.

(a) Four phylogenetically novel *Caudoviricetes* SA-vFAMs, with the boxed region illustrating the comparative genomic analysis of representative vOTUs in SA-vFAM9.

(b) ViPTree analysis demonstrates evolutionary divergence between SA-vFAM9 and ICTV reference sequences, identifying it as a novel family.

(c) Proteomic similarity network generated via vConTACT2 using Bacteroidales phages from INPHARED as references (dark grey dots).
(d) Comparison of sequence lengths between active and inactive vOTUs (Wilcoxon test).
(e) The sequencing depth distributions of the three SA-vOTUs within vFAM7.
(f) TerL-based phylogenetic tree of SA-vFAM7 and 1,032 *Ca. Heliusvirales* reference sequences. Rings from inside to outside represent: (1) the top 5 known families within *Ca. Heliusvirales*; (2) the activity of each SA-vFAM7 genome; (3) the host taxonomy of SA-vFAM7 members; and (4) the distribution of SA-vOTUs within SA-vFAM7.
(g, h) Characterization of AMG insertions within vSUBFAM7 at the HNH-TerS (g) and TerS-TerL (h) intergenic regions. Sub-panels include the distribution of insertion-carrying genomes across different genera and the functional annotations of the inserted genes.

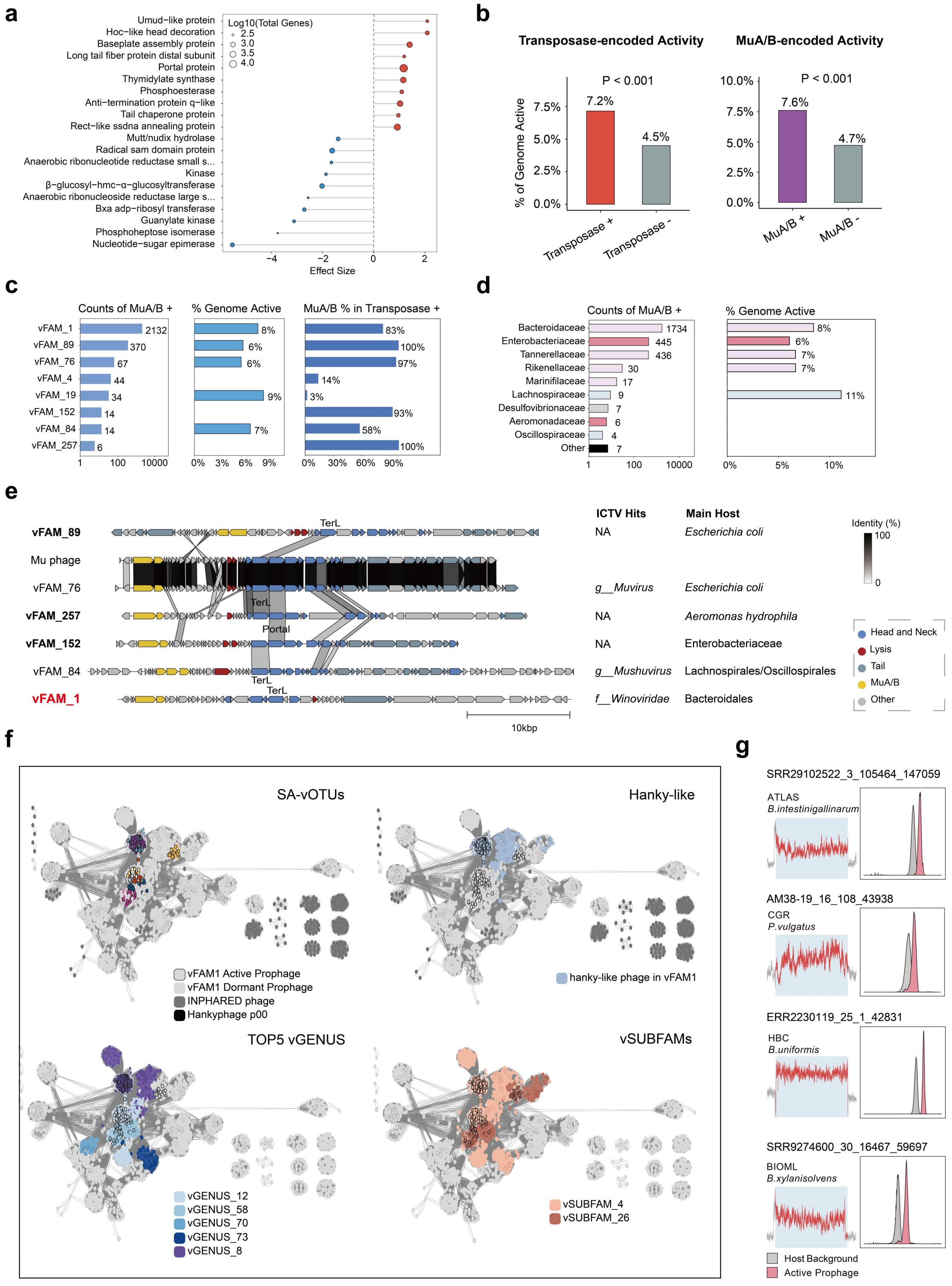

Figure S6. Genomic feature enrichment in active prophages reveals active Mu-like vFAMs.

(a) Top 10 annotation terms enriched in active and inactive prophages. Left and middle panels show the statistical significance derived from GLMM.

(b) Proportion of active phages among prophages stratified by transposase and MuA/B carriage. Statistical significance was evaluated using Fisher's exact test.

(c) Taxonomic origins and corresponding activity levels of MuA/B phages across vFAMs (left), counts of MuA/B members per vFAM (middle), and the proportion of Mu-like members among the transposase-encoding population within each vFAM (right).

(d) Host distribution of all identified MuA/B phages, detailing counts (left) and their respective proportion of active prophages (right).

(e) Genomic architecture of the six identified gut Mu-like phage vFAMs. Schematics highlight the conserved MuA/B operons, alongside common host and established ICTV taxonomic classifications (below class level).

(f) Proteomic similarity network of vFAM1 generated via vConTACT2 using Bacteroidales phages from INPHARED as references. Sub-panels sequentially highlight the topological distribution of SA-vOTUs (colored), hanky-like prophages, vSUBFAMS encompassing hanky-like members, and the vGENUS lineages containing these prophages. Active and dormant prophages are differentiated by distinct node-border outlines.

(g) Representative SA-vFAM1 sequencing depth profiles validating prophage activity.
